# Alternative splicing of a TPR domain determines mitochondrial versus plastid function of the only CLU family protein in *Marchantia polymorpha*

**DOI:** 10.64898/2026.03.13.711532

**Authors:** Maria Lozano-Quiles, Parth K. Raval, Sven B. Gould

**Affiliations:** Institute for Molecular Evolution, Heinrich-Heine-University Düsseldorf, 40225 Düsseldorf, Germany; Institute for Population Genetics, Heinrich-Heine-University Düsseldorf, 40225 Düsseldorf, Germany

**Keywords:** *Marchantia*, Alternative splicing, Chloroplasts, Mitochondria, Organelle distribution, TPR domain

## Abstract

In plant cells, the multi-domain proteins FRIENDLY and REC regulate the cellular organization, distribution and proliferation of mitochondria and plastids, respectively. Both proteins share a similar overall domain architecture and belong to the larger CLUSTERED MITOCHONDRIA (CLU) superfamily. Domains of CLU proteins have been shown to interact with translation related proteins, tRNA synthetases and even mRNA, but their exact modes of operation remain cryptic and how organelle specificity of CLU paralogs in plant cells is achieved remains unknown. We characterized the single CLU family protein of the liverwort *Marchantia polymorpha* that we demonstrate to be transcribed either with or without exon 22, which changes the configuration of the TPR domains in the C-terminus. Knockout of Mp*CLU* affects both mitochondria and plastids, and independent rescues show that the splice variant with exon 22 (MpCLU^22^) serves mitochondrial- and the one lacking exon 22 (MpCLU^spl22^) plastid biology. The CLU-C domain of the protein when expressed alone localises to the nucleus and induces a phenotype that differs in photosynthesis performance and transcriptome changes from that of the knockout of Mp*CLU*. Our results identify the C-terminal TPR motif to be responsible for organelle specificity in plants and they provide an example of how genome reformatting and gene loss can be compensated for by the alternative splicing of a single exon.

## Introduction

The origin of eukaryotes added a new type of cell to the tree of life, a cell characterized by several membrane-bound compartments that together form the endomembrane system. Eukaryotic compartments evolved during eukaryogenesis in an archaeal host, in an unknown order, and likely as a consequence of endosymbiosis (Raval et al., 2022; Richards et al., 2024). The cytosolic space allocated to and the distribution of any given compartment is no coincidence, but the result of a continuously orchestrated feedback loop (Chan and Marshall, 2010; Lloyd AC, 2013). For example, the nuclear volume is fine-tuned with respect to cell size by proteins of the lamin family, and the LINC and nuclear pore complex (Jevtić et al., 2014; Y. Wu et al., 2022). The Golgi apparatus is more dynamic, its volume being regulated by proteins such as Sec16 and Src kinases and depending on incoming versus outgoing vesicle flux (Sengupta and Linstedt, 2011). For two compartments, mitochondria and plastids, the regulation of their number, volume and distribution is a special case, because both were once free-living bacteria and they hence cannot be synthesised *de novo* from scratch.

One gene associated with intracellular mitochondrial distribution was initially identified in Dictyostelium and named *cluA* due to the clustering of mitochondria observed upon its deletion (Zhu et al., 1997). Subsequently characterized orthologues in Saccharomyces (*clu1*) (Fields SD et al., 1998), Drosophila (*clueless*) (Cox and Spradling, 2009), human (*cluH*) (Gao et al., 2014), and Arabidopsis (*friendly mitochondria*, *fmt* or *friendly*) (Logan et al., 2003) established the CLUSTERED MITOCHONDRIA (CLU) superfamily, as defined by the presence of CLU domains. This was followed by the characterisation of a CLU family protein from Arabidopsis, which was found to influence plastid number and morphology, and termed REDUCED CHLOROPLAST COVERAGE (REC) (Larkin et al., 2016). Orthologues thereof have since been analysed in the monkeyflower *Erythranthe lewisii* (termed REDUCED CAROTENOID PIGMENTATION or RCP) (Stanley et al., 2020), in tomato (SlREC) and other plants (Hu et al., 2024a), and it was suggested that these plastid regulators once originated after the gene duplication of a mitochondrion-associated CLU protein (Hu et al., 2024a; Larkin et al., 2016; Stanley et al., 2020).

Indeed, plastid-type CLU proteins (CLU^pl^) are remarkably similar to CLU proteins associated with mitochondrial biology (CLU^mt^). With few exceptions, they share the same identifiable domains, including multiple eponymous CLU domains and a series of tetratricopeptide repeats (TPR) in the C-terminus of the protein (Fig. 1a). This raises the question of how CLU paralogues within the same species (e.g. FRIENDLY vs REC in Arabidopsis) serve each organelle specifically and how the CLU^pl^ subfamily functionally diverged from the CLU superfamily. Despite the high level of sequence and domain architecture similarities, CLU^mt^ and CLU^pl^ proteins show distinct subcellular localisation and functions. CLU^mt^ protein localisation is predominantly cytosolic, but sometimes they are found to cluster around mitochondria (Cox and Spradling, 2009; El Zawily et al., 2014; Kumar et al., 2002) and to contribute to mitochondrial distribution, fission, and mitophagy (Ayabe et al., 2021; Cox and Spradling, 2009; El Zawily et al., 2014; Fields SD et al., 1998; Gao et al., 2014; Zhu et al., 1997). CLU^mt^ proteins interact with ribosomes and other translation-related proteins, and in animals have been reported to assist with the import of nuclear-encoded mitochondrial proteins (Gao et al., 2014; Ma et al., 2021; Miller-Fleming et al., 2025; Sen and Cox, 2016; Yang et al., 2022). In contrast, select CLU^pl^ proteins (e.g. At*REC1* and its homologues) show cyto-nuclear localisation and regulate the number and cellular space allocated to plastids (Larkin et al., 2016), as well as chlorophyll- and carotenoid-biosynthesis (Hu et al., 2024a; Stanley et al., 2020). The molecular details of how the CLU^mt^ and CLU^pl^ subfamilies distinguish between the two organelles, how they shift localisation from the cytosol to the nucleus or how they secure organelle homeostasis, however, remains uncertain.

**Fig. 1:**
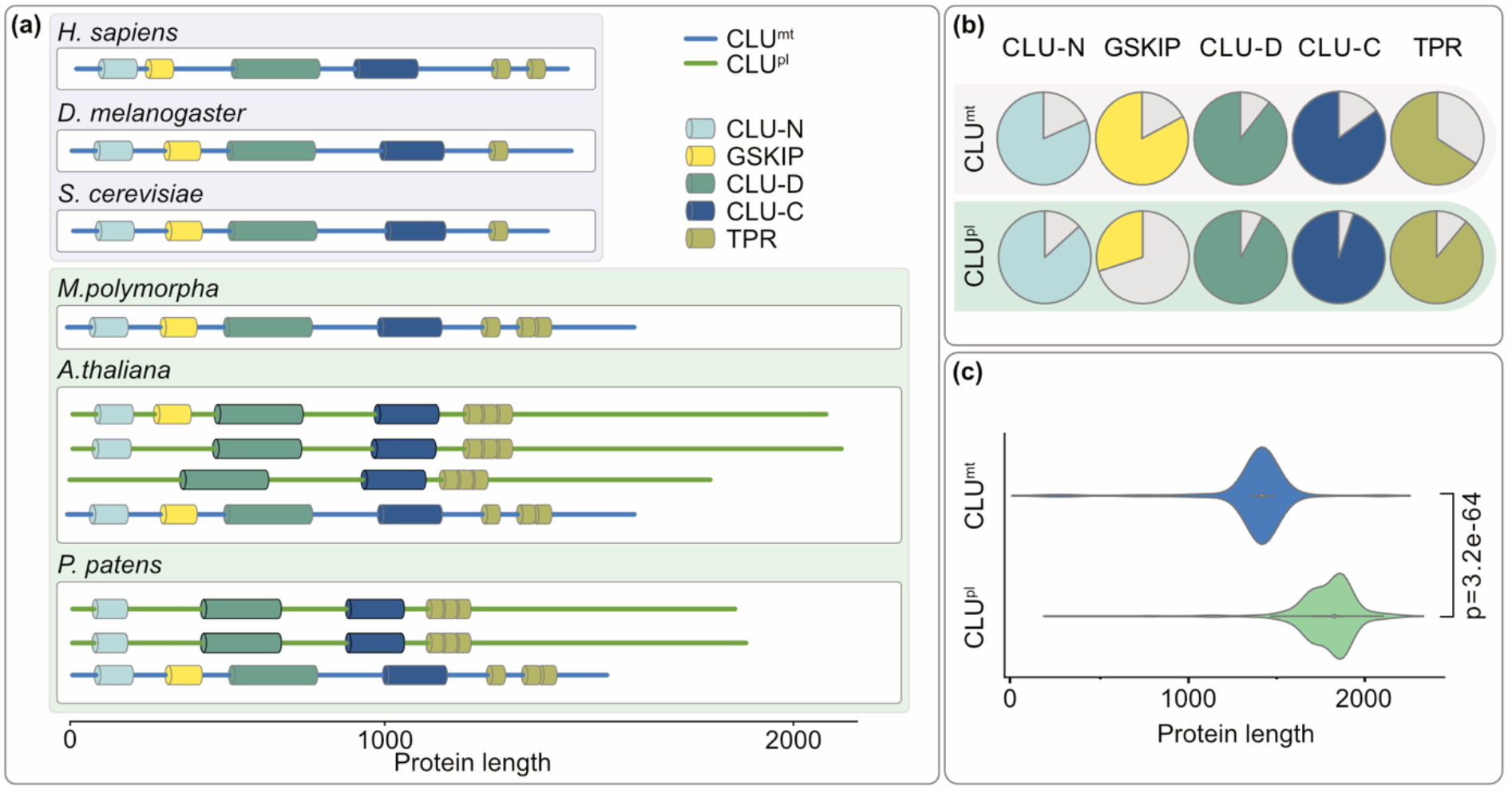
CLU^mt^ and CLU^pl^ subfamilies differ in their domain architecture and length. **(a)** Functional domain architectures based on InterPro scans for exemplary animal and plant CLU proteins. A blue backbone indicates mitochondrial-type CLU, while a green backbone indicates plastid-type CLU. TPR domains are shown as two or three individual predictions by InterProScan (SMART entry: SM00028). CLUH (KlAA0664), CLU (CG8443), CLU1 (YJM1388), MpCLU (Mp6g08800), AtREC1 (At1g01320), AtREC2 (At4g28080), AtREC3 (At1g15290), AtFMT (At3g52140), PpREC1 (Pp3c8_14980V3), PpREC2 (Pp3c24_6750V3), PpFRIENDLY (Pp3c16_5830V3). **(b)** A summary for 154 plastid- and 460 mitochondria-type CLU proteins from 133 eukaryotes with respect to the presence (coloured portion of the pie chart) or absence (light grey) of a given motif. **(c)** Lengths of CLU homologues in amino acids (p-values calculated using Wilcoxon Mann Whitney rank-sum test).

Ancestral state reconstruction of the CLU superfamily has revealed a curious distribution of homologs among bryophytes (liverworts, hornworts and mosses) (Raval et al., 2024), a group reported to encode a larger repertoire of gene families than vascular plants (Dong et al., 2025). The moss *Physcomitrium patens* encodes the canonical set of CLU paralogues that could serve both organelles, but the hornwort *Anthoceros agrestis* only two CLU^pl^ proteins, and the liverwort *Marchantia polymorpha* only a single CLU^mt^ protein. In this study, we characterized the single CLU superfamily member of *M. polymorpha* (Mp6g08800). Our data uncover (i) how the CLU^pl^ and CLU^mt^ subfamilies diverged during the time of plant terrestrialization, (ii) the domain being imported by the nucleus, and (iii) how the alternative splicing of a single Mp*CLU* gene leads to two proteoforms with distinct functional fates. The latter presents us with a unique feature from which to better understand this protein family and the overall mechanism of differential organelle volume and dispersion regulation that is so far lacking from the existing literature on both animals and plants.

## Results

### The CLU^mt^ superfamily originated during eukaryogenesis and the CLU^pl^ subfamily in the algal ancestors of land plants

A range of different proteins contribute to cellular space allocation and volume regulation of mitochondria and plastids, the two organelles of endosymbiotic origin (Larkin, 2022; Logan et al., 2003). Comparative genomics show that most of the plastid regulators originated in streptophyte algae and duplicated subsequently in land plants (Fig. S1), in line with the general trend of an increasing number of organelle-associated proteins in embryophytes (Raval et al., 2024). A global screening with InterProScan (Fig. S2) fails to identify any gene containing the eponymous clu domain across prokaryotic genomes, but finds it encoded by genomes of all eukaryotic supergroups, underscoring its role as a conserved eukaryotic signature protein. Sequence-based clustering of whole proteome models from 133 eukaryotes clearly separated CLU^mt^ and CLU^pl^ subfamilies (Fig. S3). The CLU^pl^ proteins are absent in glauco-, rhodo- and chlorophyte algae, tracing their previously suggested origins from CLU^mt^ (Larkin et al., 2016; Hu et al., 2024a) to streptophyte algae. Subsequent divergence of the two organelle-specific subfamilies is evident in predicted domain architectures (Fig. 1). The GSKIP domain, likely connected to mitochondrial metabolism and morphology (Chou et al., 2006; Loh et al., 2015; N. S. Wu et al., 2022), is gradually disappearing from the CLU^pl^ homologs (Fig. 1b). Arrangement of the TPR domains also diverged between CLU^mt^ and CLU^pl^ proteins, as well as between the CLU^mt^ proteins of plants versus animals (Fig. 1a, Fig. S4). The region following the TPR motif has been extended by approximately 500 amino acids on average in the CLU^pl^ proteins (Fig. 1c).

### *Marchantia* encodes a single *clu* gene that is alternatively spliced at exon 22

For the single Marchantia *CLU* gene (Mp6g08800), which clusters with the CLU^mt^ proteins (Fig. S3a), two gene models are annotated. They differ at exon 22, leading either to a splice variant with (MpCLU^22^) or without exon 22 (MpCLU^spl22^). The alternative splicing generates two proteoforms that differ in a stretch of only 23 amino acids immediately upstream of the TPR motif (Fig. 2a). This additional exon is enriched in proline residues, through which is it is likely to bend the peptide chains, particularly affecting the alpha-helix rich TPR motifs. As per this expectation, the structures of MpCLU^22^ and MpCLU^spl22^ are predicted to differ in the TPR motif encoding region (Fig. 2b). This is also in line with our observation that the sequence divergence of the TPR and C-terminal region helps to specifically allocate a CLU protein to either the mitochondrial or plastid subfamily (Fig. 1; Fig. S4). Furthermore, the acetylation of two lysines in At*FMT* appears to be linked to its mitochondrial specificity (El Zawily et al., 2014) and splicing leads to the loss of a lysine residue in Mp*CLU* in that very region (Fig. 2a). This does not provide molecular details on the mechanism, but narrows down the regions of the multi-domain protein that are key for the differential regulation of the two organelles.

**Fig. 2:**
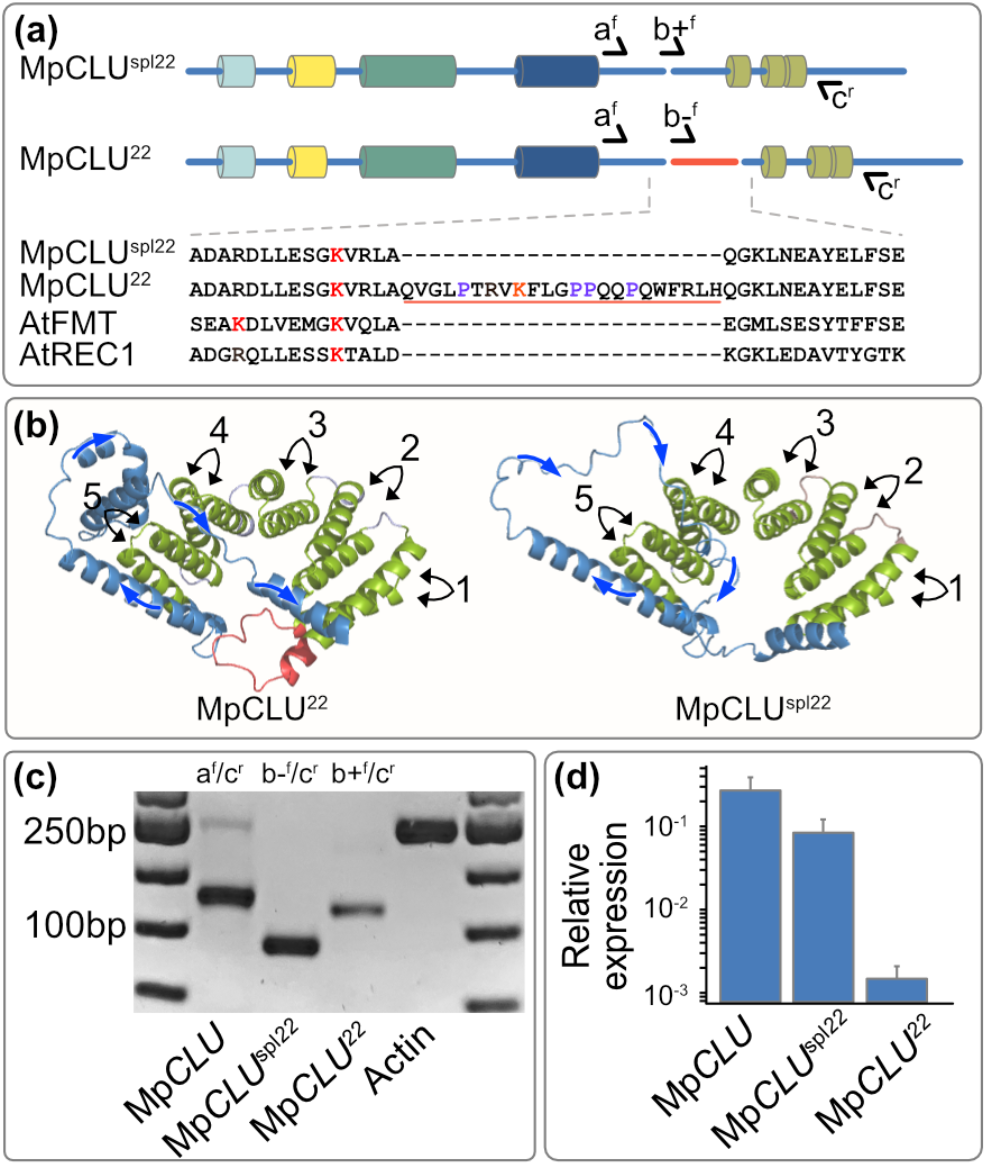
Mp*CLU* is alternatively spliced at exon 22. **(a)** Scheme of Mp*CLU* indicating the primer binding sites for specifically amplifying either Mp*CLU*^22^ or Mp*CLU*^spl22^. Exon22 in red (not at scale). Two lysine residues known to be acetylated in AtFMT and potentially corresponding residues of MpCLU are highlighted in red; proline residues present of exon 22 shown in violet. **(b)** Structural predictions of the TPR region of the two Marchantia CLU proteoforms shows that the peptide encoded by exon 22 (in red) affects the angle of an alpha helix (blue arrows) immediately after the fifth TPR domain that returns to the TPR motif. Absence of exon 22 creates a larger gap between the TPR domains. **(c)** PCRs based on complementary DNA (cDNA), indicating the primer sets used on top of each lane. **(d)** Expression levels of CLU and its splicing variants relative to that of actin (Mp6g11010), measured by qPCRs from the same cDNA as c).

To confirm the annotated gene models at the transcript level, we amplified the splicing region of interest from complementary DNA with primer pairs that amplify either one or the other splicing variant (Fig. 2a). The size differences of the products and their sequencing confirmed the expression of both Mp*CLU*^22^ and MpCLU^spl22^ (Fig. 2c). Quantitative PCRs showed that MpCLU^spl22^ is the predominant expression variant (Fig. 2d). The validation of alternative splicing, its predicted impact on a key TPR motif, and the lack of a canonical CLU^pl^ protein led us to hypothesize that alternative splicing of Mp*CLU* evolved to reinstate a CLU^pl^ protein after its loss in Marchantia.

### The CLU-C domain of Mp*CLU* localises to the nucleus and alters growth but not photosynthesis performance

Both mitochondria- and plastid-type CLU family members are multidomain proteins (Fig. 1a). CLU^mt^ proteins have been localised to the cytosol in animals and plants and found to sometimes cluster on the mitochondrial surface (Cox and Spradling, 2009; El Zawily et al., 2014; Kumar et al., 2002), while CLU^pl^ homologs have been reported to localize to the cytosol and the nucleus in several plants (Larkin et al., 2016; Stanley et al., 2020; Zhang et al., 2023). In the liverwort, the full length MpCLU^22^ was found to form small clusters in close proximity to the mitochondrial surface, whereas MpCLU^spl22^ was evenly distributed in the cytosol (Fig. 3a, S5). The CLU-N domain and the CLU-N domain in conjunction with the subsequent GSKIP domain localised to the cytosol, as did the CLU-D domain. In contrast, the CLU-C domain localised to the nucleus, as confirmed by both DAPI staining and the visible nucleoli (Fig. 3b, S6). This construct’s expression furthermore induced a growth phenotype not observed for any of the other constructs (Fig. 3b, S7). CLU-C:Citrine (referred to as CLU-C hereafter for simplicity, as with other localisation constructs) lines grew dwarfed, with a darker and disturbed thallus and thalli area and biomass were reduced by more than 80% (Fig. S7a-b). The darker shade of CLU-C lines is likely explained by the increased chlorophyll content of 50% compared to WT, along with elevated lutein, neoxanthin, violaxanthin, and beta-carotene levels (Fig. S7e-f). Despite the doubling of the chlorophyll concentration, no significant changes were measured in photosynthetic efficiency for the CLU-C lines, except for a better recovery of Y(II) and ETR after saturation of PSII (S7c-d). Multiple attempts at trying to localise the C-terminal region, including the TPR domain as well as TPR domain of MpCLU^22^ alone, failed to generate any viable lines.

**Fig. 3:**
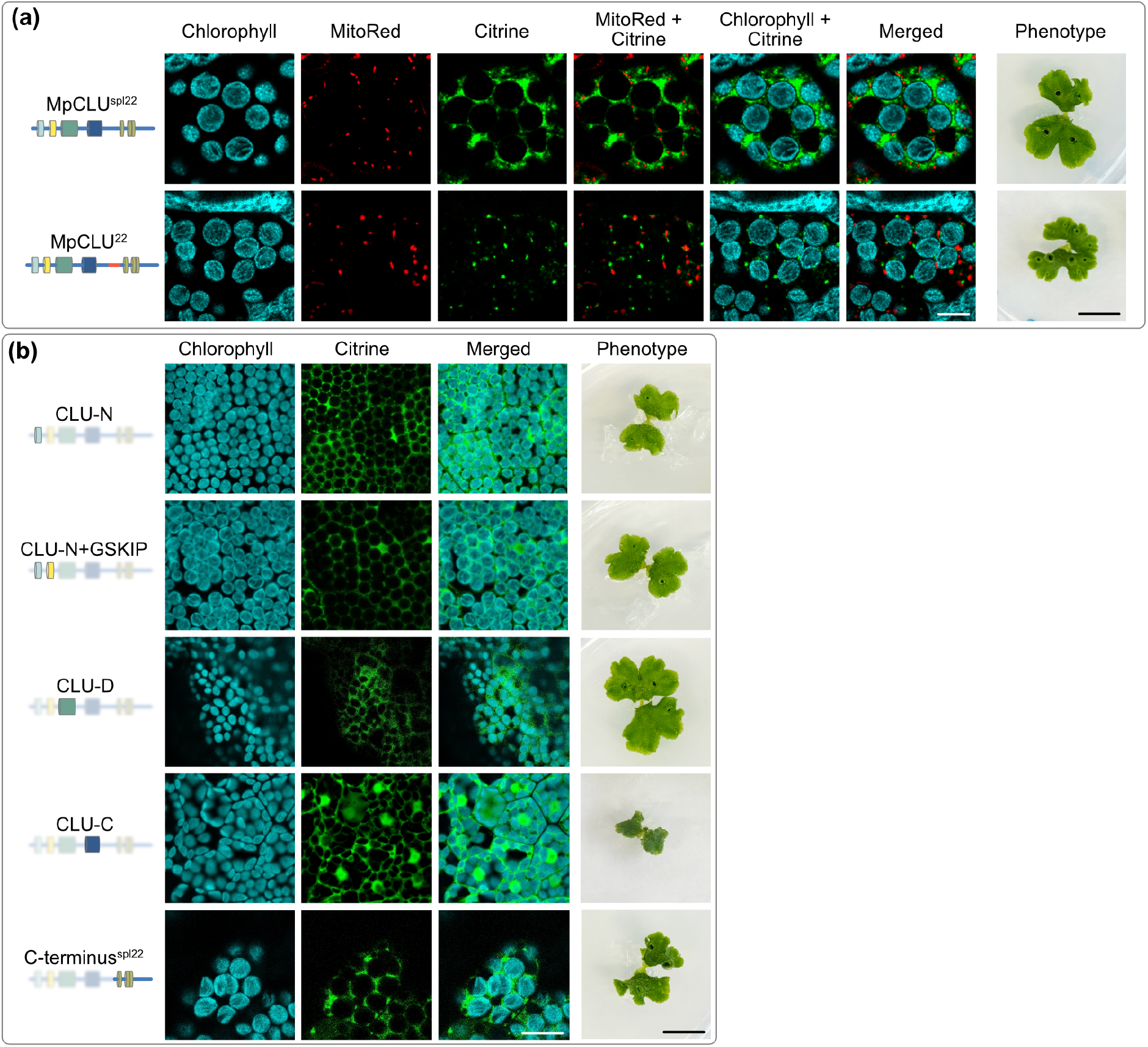
Subcellular localization of MpCLU and its individual domains fused to Citrine. Confocal laser scanning images of **(a)** the full length MpCLU^22^:Citrine and MpCLU^spl22^:Citrine **(b)** and CLU-N:Citrine, CLU-N-terminal:Citrine, CLU-D:Citrine, CLU-C:Citrine and C-terminus^spl22^:Citrine in one-day-old thalli along with representative images of 14d-old plants harbouring the respective constructs. All fusion constructs were expressed under CaMV 35S promoter in stable and isogenic transfectant plant lines. Plastids imaged through their autofluorescence (a-b) and mitochondria through MitoTracker Red (a).

### Loss of Mp*CLU* affects both mitochondria and plastids

Using an optimized CRISPR-Cas9 protocol (Sugano et al., 2018), we generated loss of function mutants for Mp*CLU* (Δ*clu*). For a detailed characterisation, we choose a mutant line in which a set of guide RNA introduced an early stop codon (Fig. S8). The Δ*clu* mutants were paler and, in comparison to WT plants, grew well below half the rate (Fig. 4a, b). The thallus size and biomass were reduced by approximately 90% (Fig. 4b) and even after six weeks, the formation of gemma cups or gametangia was rarely observed (Fig. 5). At the cellular level, the loss of Mp*CLU* on the one hand resulted in the clustering of mitochondria and on the other in 70% fewer plastids (Fig. 4c, d, S9a). The latter appeared to be compensated for by the almost doubling of organelle size (Fig. 4c, d, S10a). Hence, the total area occupied by plastids in *Δclu* cells is similar to that of WT cells (Fig. S10a-b). The number of mitochondria per cell and the area occupied by mitochondria remained rather stable across lines (Fig. S10b).

**Fig. 4:**
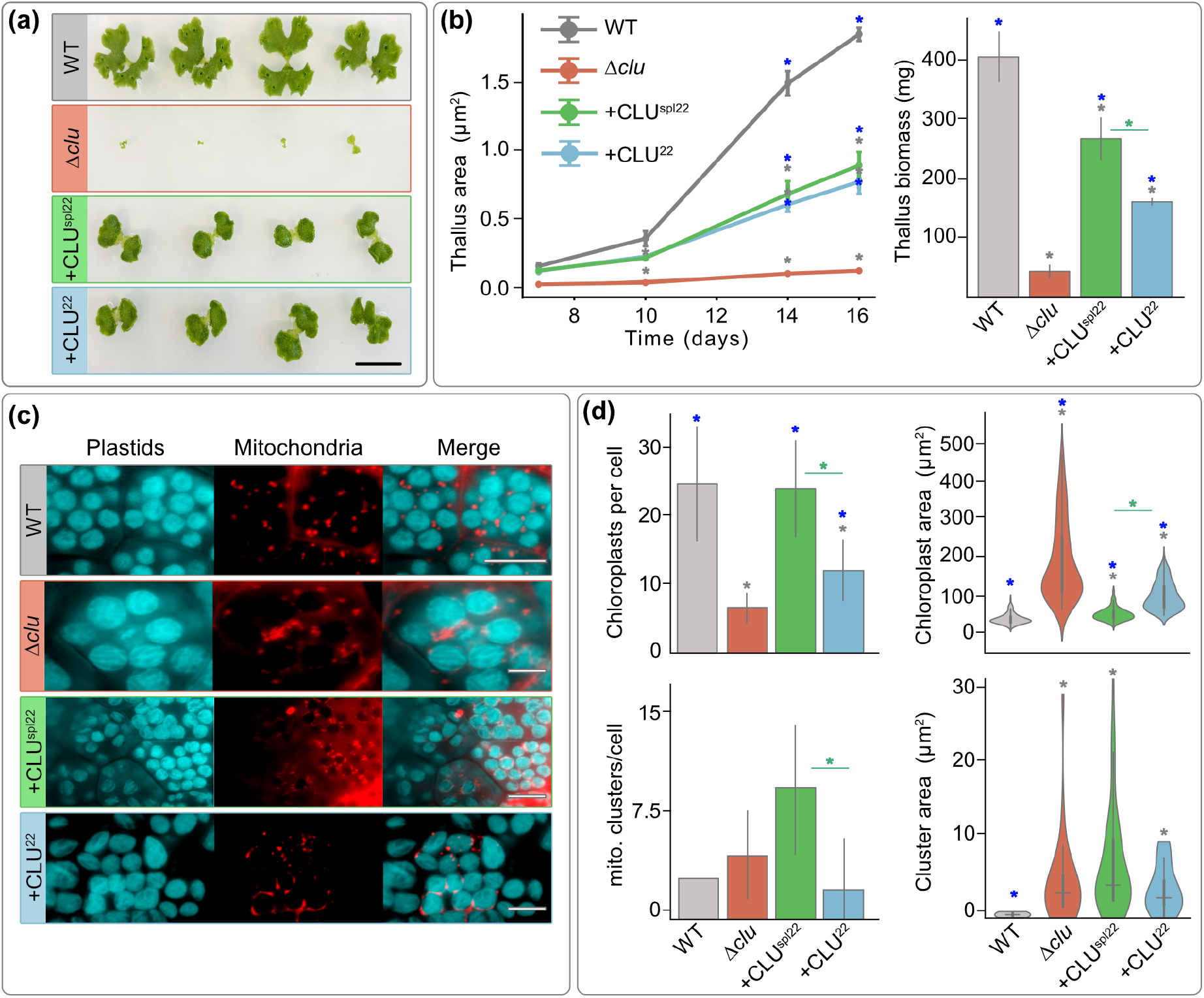
The difference in the C-terminal TPR domain is responsible for plastid versus mitochondrial phenotype rescue. **(a)** Representative images of WT, Δ*clu*, Δ*clu* + CLU^spl22^ and Δ*clu* + CLU^22^ plants grown from gemmae till 14 days (scale bar: 1 cm). **(b)** Area measurement (left) and biomass (right) of the same plants as in (a); asterisks indicate a significant difference compared to the wild-type with p-value<0.05 (n=4). **(c)** One-day-old thalli pictures of WT, Δ*clu* and Δ*clu* rescued with either CLU^spl22^ or CLU^22^. Plastids were imaged through their autofluorescence (a-b) and mitochondria through MitoTracker Red Scale bar: 10 µm. **(d)** Above, number of chloroplasts per cell and chloroplast area of WT, Δ*clu* and Δ*clu* rescued with either CLU^spl22^ or CLU^22^, one-day-old independent thalli (n=10 thalli and 15 cells per line) grown from gemmae. Below, mitochondrial cluster numbers per cell and the area occupied by mitochondrial clusters of the same plant lines, one-day-old gemmae were analysed (n=10 gemmae and 15 cells). Grey, blue and green asterisks indicate significant differences compared to the WT, Δ*clu*, and those between Δ*clu* + CLU^spl22^ and Δ*clu* + CLU^22^, respectively (p-value<0.05; two-sided Mann–Whitney U test).

**Fig. 5:**
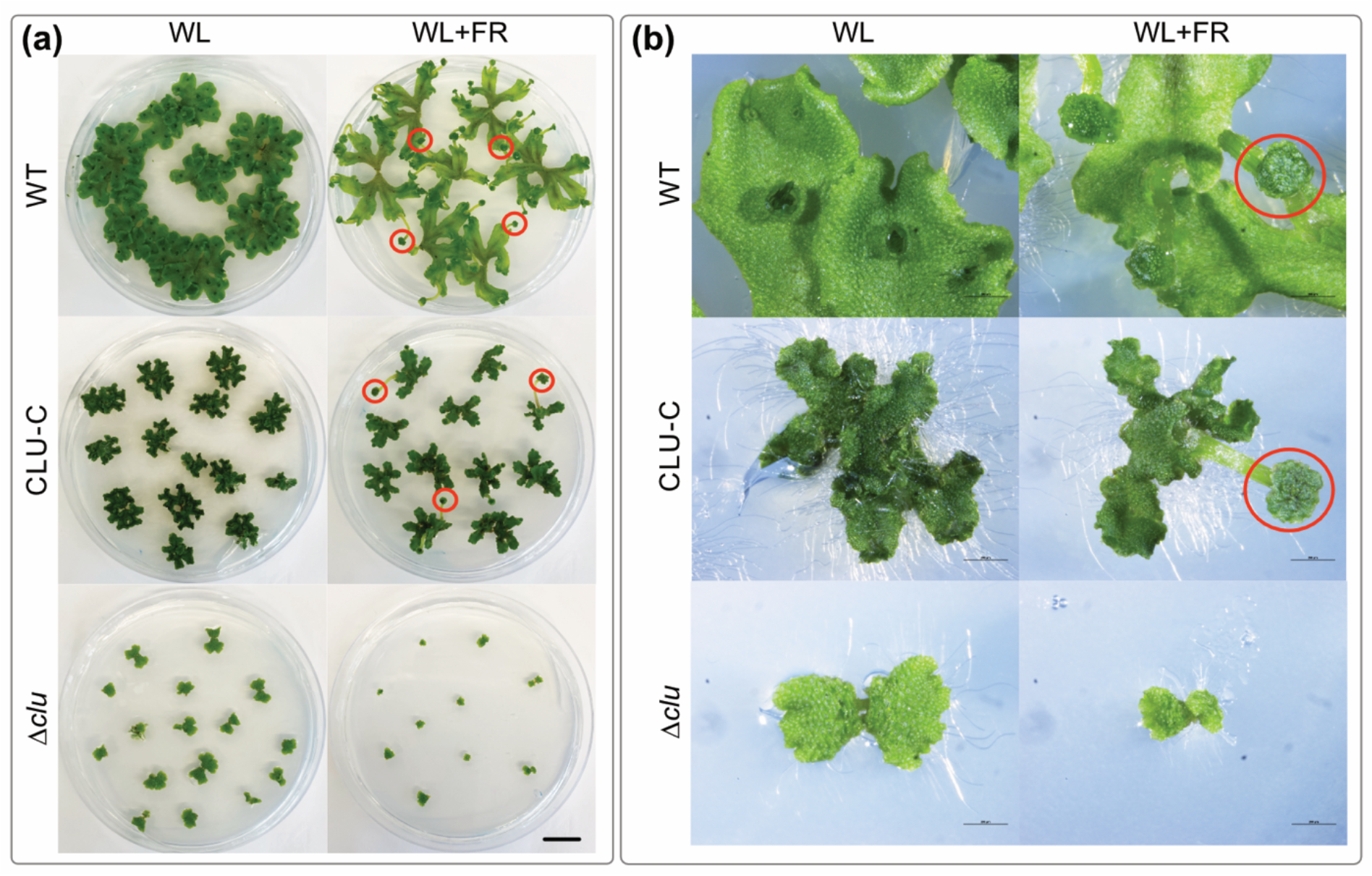
*Δclu* and CLU-C:Citrine plants show growth defects and a delay in the formation of reproductive organ. When grown under far-red (FR) light conditions, vegetative-to-reproductive state shift is induced, which involves the formation of gametangiophore structures (red circles). **(a)** 16-day-old representative plants of WT, *Δclu* and CLU-C:Citrine lines incubated under white light (WL) or WL supplemented with FR light. Scale bar: 1 cm. **(b)** Zoom in pictures of plants shown in (a) taken with binocular Nikon SMZ18. Scale bar: 0,2 cm

RNA-seq from two-week-old thalli showed that *Δclu* plants downregulated 300 genes involved in cell wall remodelling, abscisic acid biosynthesis and signalling, detoxification, anti-desiccation response, solute/water transport, protein degradation and carbon metabolism, early light inducible proteins (ELIPs), and metabolism-related genes such as the TCA cycle enzyme Mp*OGDH* (Mp4g05670; an ortholog of α-ketoglutarate dehydrogenase) suggesting changes in mitochondrial metabolic pathways (Fig. S11-12). In contrast, CLU-C:Citrine plants downregulated 240 genes mostly pertaining to cell wall defence and cell surface remodelling, Cu/Ca homeostasis, defence signalling related to salicylic acid, and also ELIPs (Fig. S11) However, Mp*AOX1* (Mp6g09630; a canonical marker for mitochondrial damage; El Zawily et al., 2014) was significantly upregulated in CLU-C:Citrine lines (Fig. S12), hinting that an excess of CLU may also affect mitochondrial function.

We complemented *Δclu* by a transfection with either the MpCLU^22^ or the MpCLU^spl22^ splice variants and first validated the expression of respective splice variants in two independent rescue lines per variant (Fig. S13a) in order to evaluate the effect of the presence and absence of exon 22. The presence of MpCLU^spl22^ in the *Δclu* background significantly improved the growth rate of the plants from all lines (Fig. S13b). Of the two independent rescue lines per variant, one of each (that had a comparable expression of the complemented constructs in comparison to the other) was chosen for further characterization. Both these lines reached about 40% of the WT thallus area and 70% biomass after 14 days of growth (Fig. 4a, b). While the mitochondria continued to cluster (Fig. 4c, d, S9a), the plastid phenotype was fully restored (Fig. 4c, d, S10, S13c). The complementation of *Δclu* with MpCLU^22^ also rescued the growth similar to MpCLU^spl22^: after 14 days, the rescue lines reached about 40% of the WT thallus size and 40% of its biomass (Fig. 4a, b). Yet, while the mitochondria showed a WT-like distribution and no obvious clustering (Fig. 4c; S9a), the number of chloroplasts was lower than WT and their combined area occupied remained similar to that *Δclu* (Fig. 4c, d, S10b). The results show that complementation with MpCLU^spl22^ preferentially restores plastid morphology, whereas that with MpCLU^22^ preferentially restores mitochondrial distribution.

### Loss of MpCLU alters pigment profiles and is associated with changes in photosynthetic performance

To characterize the photosynthesis performance, we screened the plant lines using pulse amplitude modulation (PAM) imaging. Consistent with a lower level of chlorophyll in the Δ*clu* plants, we observed a reduction in the maximum quantum efficiency of their PSII photochemistry (F_v_/F_m_), hinting at a compromised function of PSII (Fig. 6a, S14). The quantum yield of photochemical energy in PSII (PSII operating efficiency; Y(II), the fraction of open PSII reaction centres (qL), and the electron transport rate (ETR) were lower in *clu*, while the capacity for non-photochemical quenching (NPQ) remained comparable to WT (Fig. 6b, S15). After saturating the photosystems with a pulse of 800 µmol photons m^-2^ s^-1^, and when Y(II) equalled 0, a pulse of 2 µmol photons m^-2^ s^-1^ was applied to measure recovery. *Δclu* did not recover as fast as WT in terms of Y(II), YNPQ(II) qL, and ETR, hinting again at a compromised PSII (Fig. 6b, S13). In the complementation lines, MpCLU^spl22^ and MpCLU^22^, the values were restored close to WT levels, both under normal and saturating conditions, with neither line differing significantly from WT (Fig. 6b, S15-16). In line with photosynthetic performance and pale thalli, Δ*clu* plants showed a significant decline in key pigments such as chlorophyll a, chlorophyll b, lutein, neoxanthin, violaxanthin, and beta-carotene (Fig. 6c). Both proteoforms increased pigment levels to a varying extent, with chlorophyll a, chlorophyll b and neoxanthin levels being fully restored only by MpCLU^spl22^ (Fig. 6c). Altogether, these results indicate that the loss of MpCLU compromises both PSII activity and pigment accumulation, resulting in impaired photosynthetic performance. The complementation of *Δclu* suggests a stronger contribution of MpCLU^spl22^ in sustaining pigment composition, although both proteoforms seem to restore photosynthetic function to a similar degree.

**Fig. 6:**
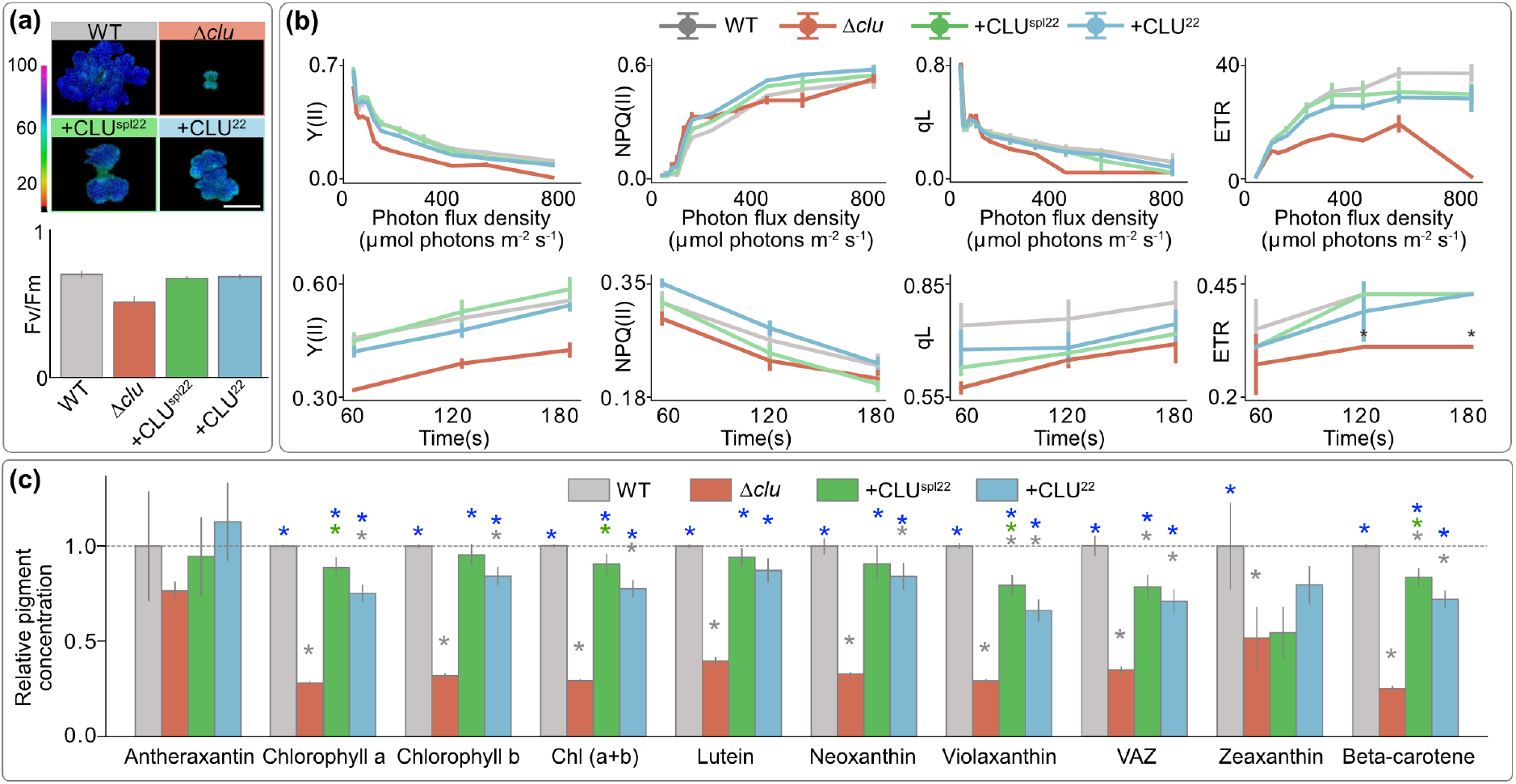
Photosynthesis and pigment concentration changes in Δ*clu* and complementation lines. **(a)** Representative images and F_v_/F_m_ values of 14-day-old plants of WT, Δ*clu*, Δ*clu* + CLU^spl22^ and Δ*clu* + CLU^22^ lines (n = 5 plants per line). Scale bar: 1cm. **(b)** Quantification of fluorescence parameters under increasing photon flux density (top): Y(II) as PSII yield, Y(NPQ) indicating regulated heat loss, qL as the fraction of open PSII reaction centres, and electron transport rates (ETR) of plants identical to those listed in a). The same parameters measured with a pulse of 2 µmol photons m^-2^ s^-1^ after PSII is saturated by a pulse of 800 µmol photons m^-2^ s^-1^. **(c)** Fold changes in pigment concentrations relative to the WT. Grey, blue and green asterisks indicate significant differences compared to the wild-type, Δ*clu*, and those between Δ*clu* + CLU^spl22^ and Δ*clu* + CLU^22^, respectively (p-value<0.05; independent two-sample Welch’s t-tests, n= three independently grown thalli mass per line).

## Discussion

We find that the alternative splicing of a single exon in the TPR region of the only Marchantia CLU protein is associated with mitochondrial versus plastid function. The divergence in domain architecture between CLU^mt^ and CLU^pl^ is key to understanding how CLU proteins evolved organelle-specific functions. CLU orthologs contain five predicted domains that offer insights into their specialization. At the N-terminus, the CLU-N domain mediates self-interaction (Ayabe et al., 2021), while the GSK3B interacting protein domain (GSKIP) – more prevalent in CLU^mt^ than CLU^pl^ – is part of an A-kinase anchoring protein complex that binds GSK3β, a kinase involved in microtubule dynamics, cell-cycle progression, metabolism, and transcription factors (Wu et al., 2022). The function of the CLU-D domain remains uncharacterized, while CLU-C is a domain through which the Arabidopsis CLU^mt^ homolog FMT interacts with translation-related proteins such as eIFiso4G1, which is known to increase the translation of chloroplast proteins (Ayabe et al., 2021; Lellis et al., 2019). At the C-terminus, the tetratricopeptide repeats (TPR) domain appears to be crucial for CLU functions, as the deletion of either the TPR domain or CLU-C prevents CLU^mt^ from rescuing the loss-of-function phenotype in Drosophila (Sen et al., 2015).

The acetylation of two lysine residues in proximity to the TPR domain of the Arabidopsis CLU protein FMT is essential for its function, as mutating either site prevents molecular complementation (El Zawily et al., 2014). We speculate that this is associated with the need for a functional motif this region represents for outer mitochondrial membrane binding, which at the same time acts as a switch, because lysine acetylation can regulate the binding of peripheral proteins to membranes by masking the positively charges residues that otherwise interact with the negatively charged bilayer (Okada et al., 2021). Acetylation of K1022 and K1029 of AtFMT regulates its activity (El Zawily et al., 2014) and in the homologues region of Marchantia, two lysine residues are only present in MpCLU^22^, the very variant we find associated with mitochondria. Notably, our phylogenetic analysis of CLU protein clusters and protein domains identify the organization of TPR motifs in the C-terminus to differ between the CLU^pl^ and CLU^mt^ isoforms across species (Fig. 2; S4). This structural variation between CLU^pl^ and CLU^mt^ is also evident in the Mp*CLU* splicing variants. It contributes to their functional divergence, likely by either providing an outer mitochondrial membrane binding region or not.

Consistent with a broader role of CLU-family proteins in both organelle’s coordination the loss of AtFMT (a CLU^mt^) suppresses the chloroplast proteostasis defects of *clp1* mutants through a compensatory mechanism involving REC1 and REC2 (both CLU^pl^) (Kim J et al., 2026). The knock-out of the single CLU homolog in *Marchantia* combines effects on mitochondria and chloroplasts, underscoring the general dual-function of the protein in the liverwort. In contrast to what was described for the angiosperm, however, the loss of Mp*CLU* leads to fewer and larger chloroplasts in the bryophyte. Complementation with MpCLU^spl22^ rescues the plastid phenotype but does not mitigate the clustering of mitochondria, while the expression of MpCLU^22^ reduces mitochondrial clustering without restoring the chloroplast phenotype. This is further evidence for a splicing-dependent, functional separation of the two MpCLU proteoforms through a simple change in the C-terminus associated with the TPR and an associated, acetylatable region (Fig. 2a) It resembles similar cases, in which alternative splicing generates functionally distinct proteins from a single gene (Kanazawa et al., 2016; Zhang and Mount, 2009). Alternative splicing of Mp*CLU* was selected for as a solution to compensate for the evolutionary reduction to a single *CLU* gene, while still needing to serve two organelles of endosymbiotic origin.

It remains uncertain how the CLU protein family measures and or communicates organelle volume and distribution throughout the cell, but their localisations are likely to play a role. Full length MpCLU^spl22^ localizes to the cytosol and is distributed evenly, consistent with previously reported CLU^mt^ homolog localization (Cox and Spradling, 2009; El Zawily et al., 2014; Kumar et al., 2002), whereas the full-length MpCLU^22^ displayed a punctuated cytosolic distribution, frequently associated with mitochondria. Hence, the exon-dependent difference in the ability to rescue either the mitochondrial or chloroplast phenotype appears to be accompanied by exon-dependent difference in localization at least for MpCLU^22^. While this alone cannot provide mechanistic insights, it does identify the critical domain. Considering that the N-terminal domains (CLU-N and CLU-N + GSKIP), as well as the C-terminus including the TPR domain localize to the cytosol, it leaves only the CLU-C domain that could potentially confer a nuclear localisation (e.g. to Arabidopsis CLU^pl^ homologue). The nuclear localisation of CLU-C is accompanied by phenotypes such as dark-green thalli, an increased carotenoid level, and an upregulation of Mp*WRKY7* (Mp3g17660), which belongs to a transcription factor family implicated in carotenoid biosynthesis (Diao et al., 2023) and Mp*AOX1* (Mp6g09630; a canonical marker for mitochondrial damage) was also upregulated in CLU-C plants (Fig. S11-12), further indicating that the expression of this isolated domain alone can affect expression of genes involved in plastid and mitochondrial processes. Together, we suggest that while the TPR motif is responsible for organelle specificity, the CLU-C domain may provide further insights into the context-dependent nuclear localization and altering gene expression.

The growth deficit of *Δclu* plants observed in this study is accompanied by a substantial reduction in their photosynthetic performance, and lower chlorophyll and carotenoid levels. PSII is significantly affected, reflected by a lower F_v_/F_m_, qL and ETR, together with an elevated YNPQ(II), indicating greater dissipation of excitation energy as heat. The chloroplasts seem to respond with a higher level of photoprotective mechanisms. The combined photosynthetic and pigment accumulation phenotype of *Δclu* plants resemble those reported for *fmt* (Arabidopsis) and *slrec* (Tomato) mutants, respectively (El Zawily et al., 2014; Hu et al., 2024b). Complementation of *Δclu* with MpCLU^spl22^ or MpCLU^22^ partially restores thallus area and biomass, indicating that both isoforms can recover key physiological functions. Although both MpCLU proteoforms restored photosynthetic performance to a similar extent, MpCLU^spl22^ more effectively recovered chlorophyll a and b, violaxanthin and beta-carotene than MpCLU^22^, that showed a more limited pigment recovery but a stronger recovery of zeaxanthin (Fig. 6) This difference suggests that although both proteoforms support the core photosynthetic machinery, MpCLU^spl22^ may be more effective in coordinating pigment biosynthesis or stability, while MpCLU^22^ may regulate stress-responsive pathways, pointing toward a functional divergence between the two isoforms or a differential contribution. These findings point to distinct, yet complementary roles for the two proteoforms in chloroplast function and photoprotection in an agreement with inter-organellar compensation of the Arabidopsis FMT (Kim et al.2026), although the underlying mechanisms remain unclear.

In summary, our findings demonstrate that in Marchantia a single *CLU* gene participates in the orchestration of both mitochondrial and plastid homeostasis through the alternative splicing of exon 22 – a strategy adopted to maintain regulatory complexity and following bryophyte genome reformatting. We identify changes in the C-terminal region of CLU proteins to be responsible for mediating mitochondrial versus plastid function after gene duplication of a mitochondrial CLU protein in a streptophyte algal ancestor of land plants. In particular the TPR motif is of importance and in CLU^mt^ proteins likely involved in binding the outer mitochondrial membrane if not acetylated. CLU proteins are a critical and conserved component of eukaryotic biology and their C-terminal region can now serve as a prime target for future studies to uncover the molecular details of how the protein family exactly contributes to the regulation of mitochondrial and chloroplast distribution and the upkeep of a functional organelle to cell volume ratio.

## Materials and methods

### Comparative genomics

We traced evolutionary history of organelle distribution control protein using a database of 157 Archaeplastidal and 10 outgroup eukaryotic genomes (species names and taxonomy summarized in Table S1A). The whole proteome models were clustered using OrthoFinder version 2.5.4 (Emms and Kelly, 2019). The resulting protein clusters were annotated based on the presence of previously characterized proteins involved in organelle volume distribution (Larkin, 2022). Homologues identified as CLU^mt^ and CLU^pl^ (based on the presence of Arabidopsis RECs and FRIENDLY in the same cluster) were further analysed for their domain architecture using InterProScan (Larkin, 2022; Paysan-Lafosse et al., 2022) and to predict their protein’s structure using AlphaFold (Jumper et al., 2021). The two protein family clusters were visualized in Cytoscape, separately from each other, using sequence similarities (Buchfink et al., 2014) of each protein with every other protein in the respective orthogroup (CLU^mt^ and CLU^pl^).

### Growth conditions

*Marchantia polymorpha* (Tak-1 ecotype) was cultivated on half-strength Gamborǵs B5 medium (GVA) (pH 5,8) containing 1% agar under continuous full light spectrum (450-700nm) 70 μmol m^-2^ s^-1^ at 22°C. Unless otherwise stated, all plant lines were grown under these conditions. Reproductive state was induced by cultivating Marchantia with the same light conditions supplemented with far-red light (700-750nm) 40 μmol m^-2^ s^-1^ at 22°C.

### Cloning and mutant plants generation

MpCLU^spl22^ (Mp6g08800.1, alias Mapoly0060s0041), MpCLU^22^ (Mp6g08800.2, alias Mapoly0060s0041), and individual domains were amplified from cDNA and cloned into the entry vector pDONR221 (Invitrogen) by Gateway BP reactions set as per manufactureŕs protocol. For localization studies, CLU-N, CLU-N+GSKIP and CLU-C were cloned into the N-terminal side of the fluorescent tag Citrine on pMpGWB105, and MpCLU^spl22^, MpCLU^22^ and C-terminal constructs were cloned into the C-terminal side of Citrine on pMpGWB106 via Gateway LR reactions following manufacturer’s protocol. All vectors used for this assay contained the constitutive promoter 35S. For loss-of-function analyses, CRISPR/Cas9-based genome editing of Mp*CLU* was performed by targeting the first exon of MpCLU. The oligonucleotides containing the target sequences were synthesized and cloned into pMpGE_En03 (Sugano et al., 2018). Generated entry clones were recombined into the expression vector pMpGE010 by Gateway LR Clonase (Thermo Fisher) as per manufacturer’s instruction. For complementation of Δclu, the CDS of Mp*CLU*^spl22^ and Mp*CLU*^22^ were cloned into pDONR221 (Invitrogen) and recombined into pMpGWB301 using Gateway LR Clonase (Thermo Fisher).

All expression vectors were transformed into *Agrobacterium tumefaciens* (GV301 without pSoup) by electroporation (Bio-Rad GenePulser Xcell, 1.44 kV). Subsequent plant transformations were carried out as previously described (Raval et al., 2025). The resulting transformants were grown and selected on half-strength Gamborg’s B5 media containing 1% phyto-agar supplemented with Hygromycin (10 μg/mL) and Cefotaxime (100 μg/mL) for localization and loss-of-function constructs, and 0.5 μM chlorsulfuron to screen complementation constructs. Gemmae from multiple explants were taken forward to the next generation and grown until they produced gemmae, which were further screened. To genotype Marchantia plant lines, 100 mg of thalli from individual plants were used to extract gDNA that was used as a template for polymerase chain reactions.

For the full-length splicing variants and individual domains with Citrine, as well as the loss of function lines and rescue, gemmae from one transfectant line per genotype were analysed, and each gemmae and a plant grown from it was considered as an independent biological replicate.

Gemmae from expression-positive plant lines were grown for 24h and placed in 30 μL of water on a slide and covered with a coverslip. Protoplasts were prepared as previously described (Raval et al., 2025). Marchantia one-day-old thalli and protoplasts were stained with 500 nM MitoTracker^TM^ CMXRos following manufacturer’s instructions. The protoplast or cell suspension were incubated with 500 nM dye for 20 min at room temperature (RT), washed three times by centrifuging the cells at 2,000g and protoplasts at 100g for 5 min to visualize mitochondria.

### Fluorescence imaging analysis

Analyses of Citrine, chlorophyll fluorescence and mitochondria were performed according to our previous studies (Raval et al., 2025). Briefly, Nikon Eclipse Ti Imaging platform and confocal laser scanning microscopy (SP8X system, Leica Microsystems, Wetzlar, Germany) were used with the settings previously described (Raval et al., 2025). Autofluorescence from chloroplasts was filtered out by the time-gated method (Kodama, 2016). All localization analyses and microscopy images processing were conducted in ImageJ (Fiji) with the BIOP JACoP plugin (auto thresholds).

### Cell quantification and analyses

Chloroplasts and mitochondria from gemmae were traced and measured to analyse number and area using Analyse particles plug-in in ImageJ (1.54f), as well as cell area. Mitochondrial clustering was calculated as the sum of all the areas of mitochondria that had no physical separation between them. Chloroplast coverage was calculated by dividing the total chloroplast area of a cell by the total area of that cell.

### Pigments content analysis

For pigment quantification, 50 mg of 14 days old gemmalings were used with three biological replicates per genotype. Tissue was grinded with 1mL acetone and 0.013g magnesium sulfate heptahydrate with a pestle till the sample is consistent. The mixture was filtered out with miracloth paper and stood for 15 min. Samples were centrifuged at 2000 x g speed and the supernatant stood for 30 min at 4°C and stored at -20°C until analysis. The pigment extracts were filtered (0.2 μm pore size) and then used for reversed-phase HPLC as previously described (Farber et al., 1997).

### Measurement of chlorophyll fluorescence

An IMAGING-PAM (M-Series; Walz) was used to assess the photosynthetic parameters in PSII. Pulse amplitude-modulated measuring light (<1 μmol m^-2^ s^-1^) was applied to determine minimum fluorescence (F_o_), and a subsequent saturating pulse of 3000 μmol m^-2^ s^-1^ was used to evaluate maximum fluorescence (F_m_ and F_m_’), and F’ is the fluorescence emission. Photosynthetic parameters were calculated using the following formulas: F_v_/F_m_ = (F_m_-F_o_)/F_m_’, Y(II)= (F_m_ʹ – Fʹ)/F_m_ʹ, NPQ = (F_m_ – F_m_ʹ)/F_m_ʹ, and qP = (F_m_ʹ – Fʹ)/ (F_m_ʹ – F_o_). During analysis with Imaging-PAM, pulse-modulated excitation and saturation pulses were achieved with a blue light-emitting diode lamp with a peak emission of 450 nm. Images of the photosynthetic parameters were displayed with the help of a false-color code ranging from black (0) to red, yellow, green, blue, and pink (100).

### RNA extraction, qPCR and sequencing

RNA was extracted using the RNeasy Plant Mini kit (Qiagen). Starting tissue was 14-day-old thalli, excluding gemmae cups, and grown under conditions described earlier. 500 ng of DNase-treated RNA was used as a template for preparing cDNA with iScript™ cDNA Synthesis kit (Bio-Rad) following manufactureŕs instructions. qPCR was performed using GoTaq® qPCR and RT-qPCR Systems kit (Promega). Reaction conditions comprised initial denaturation at 95°C for 5 min followed by 40 cycles of 95°C for 15 seconds (denaturation) and 60°C for 1 minute (annealing, extension and fluorescence reading).

For RNA-sequencing, three biological replicates were prepared for each genotype and mRNA isolated via their poly(A) tail. RNA integrity and concentration were assessed using a NanoDrop spectrophotometer. Libraries were prepared by Eurofins Genomics (Ebersberg, Germany). Sequencing was performed on an Illumina NovaSeq 6000 platform, generating paired-end reads (2 × 150 bp), with an average of ∼30 million read pairs per sample. Sequencing data was quality checked and analyzed as described previously (Frangedakis et al., 2024). Briefly, high-quality reads were retained using FASTQC and TrimGalore, aligned to *M. polymorpha* genome (v.7.1) using Kallisto (Bowman et al., 2017; Bray et al., 2016) and DEG were obtained through DESeq2 (Love et al., 2014).

## Data availability

Supplementary figures and tables are available with this submission. Transcriptome data are available in the NCBI Sequence Read Archive: PRJNA1435946. Supplementary data and figure source data are available on Zenodo (10.5281/zenodo.22704886).

## Author contributions

MLQ: conceptualization, supervision; experimental design, investigation, data curation, formal analysis, validation and visualization, writing original draft, review and editing. PKR: conceptualization, supervision, experimental design, formal analysis, visualization, writing original draft, review and editing. SBG: conceptualization; project administration; supervision; experimental design; funding and resource acquisition; data visualization; writing original draft, review and editing.

## Acknowledgements

We acknowledge support from the high-performance computing cluster (HILBERT) and Center for Advanced Imaging (CAi) at the Heinrich Heine University Düsseldorf and thank Michael Knopp, Michelle Kipp, Lennart Schwarz, Sina Kruse, Daniel Wasim Djamriani, and Margarete Stracke for their help. We thank Nora Gutsche and Sabine Zachgo of the University of Osnabrück, and Hirofumi Nakagami and Katharina Kramer of the MPI Cologne for their support with setting up *Marchantia*. We thank Peter Jahns (HHU Düsseldorf) for his help with pigment isolation and photosynthesis performance analysis. MLQ and PKR are grateful to William F. Martin (HHU Düsseldorf) for financial support.

