## Supplementary data for "Alternative splicing of a TPR domain determines mitochondrial versus plastid function of the only CLU family protein in *Marchantia polymorpha*"

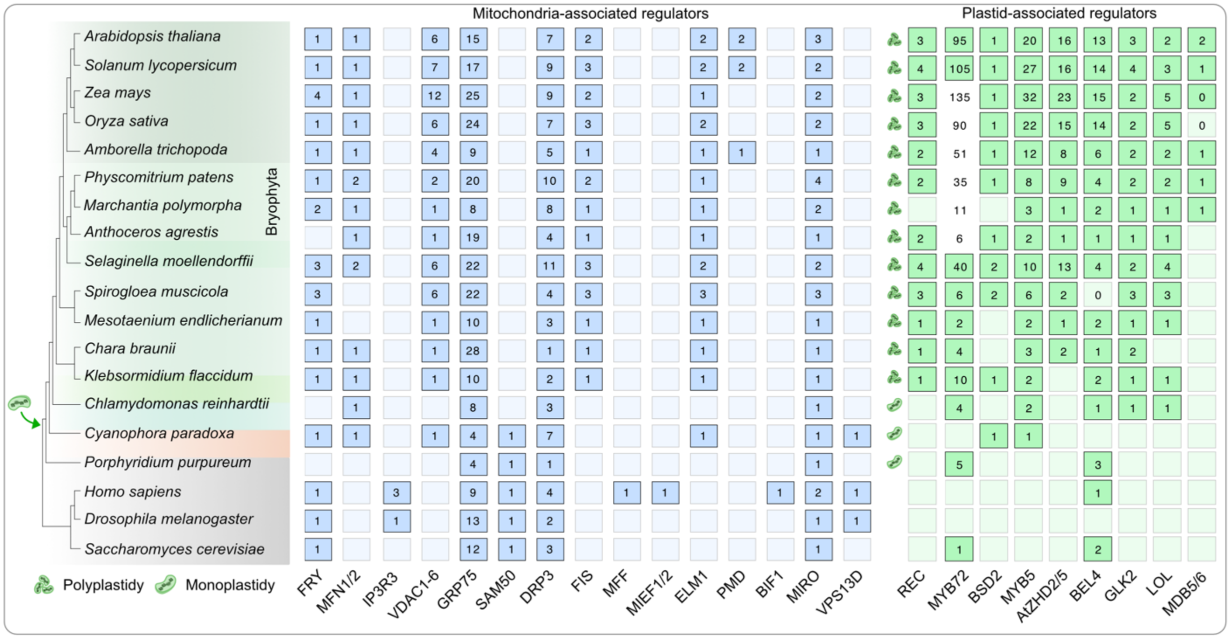


**Fig. S1: Phylogenetic distribution of proteins associated with mitochondrial and plastidial volume and distribution management.** Copy number of known organelle volume and distribution control proteins across eukaryotes reveal an early emergence of mitochondrial regulator proteins and the chloroplastidic origins of some plastid regulators, along with the increase in plastid copy number per cell (polyplastidy) in embryophytes (Arimura, 2018; Larkin, 2022).

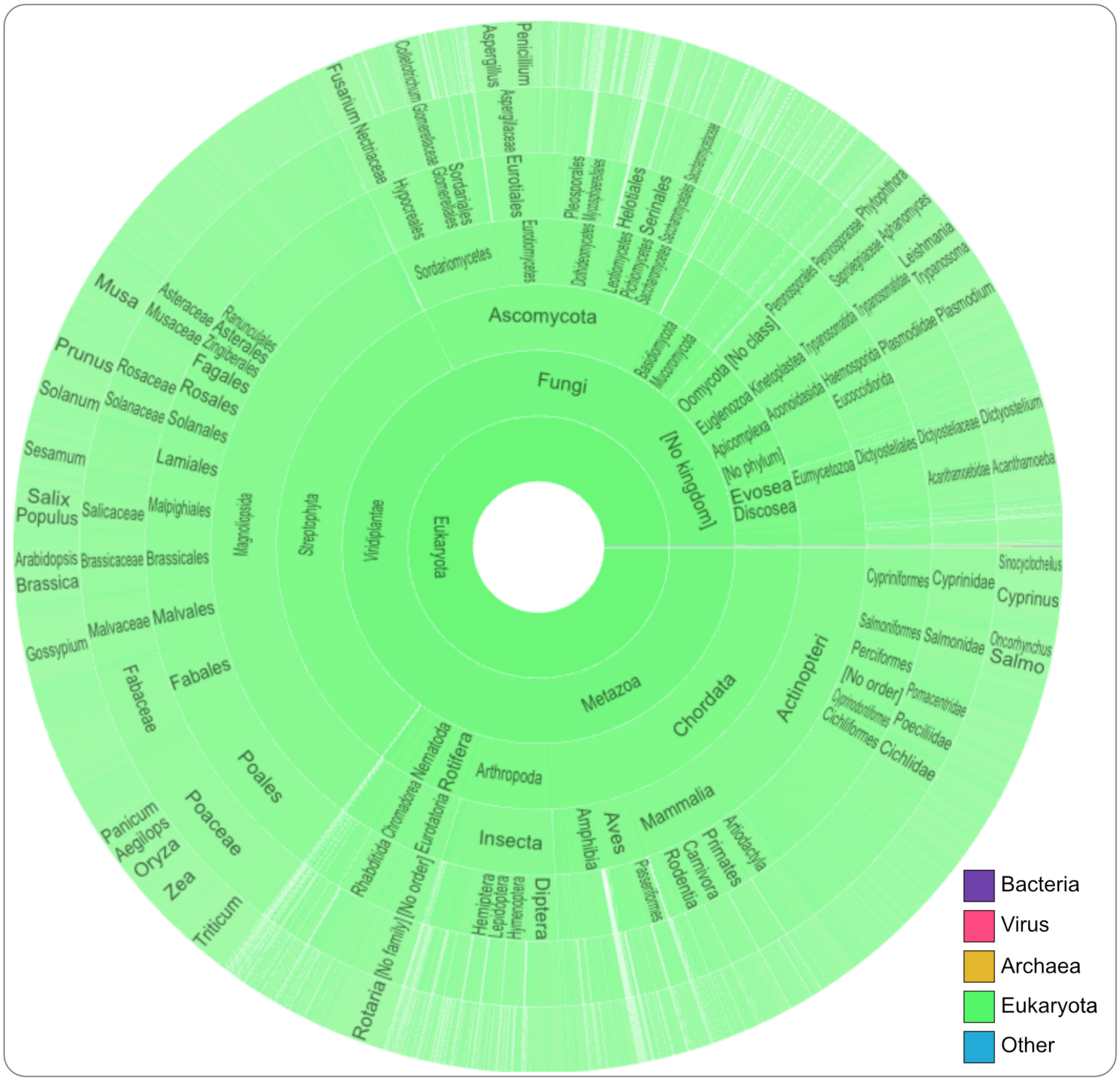


**Fig. S2: Distribution of the CLU domain across species.** The CLU domain (InterPro ID IPR025697) was exclusively found in eukaryotic genomes, suggesting it is absent from prokaryotes (as per InterPro database of August 2026, on https://www.ebi.ac.uk/interpro/).

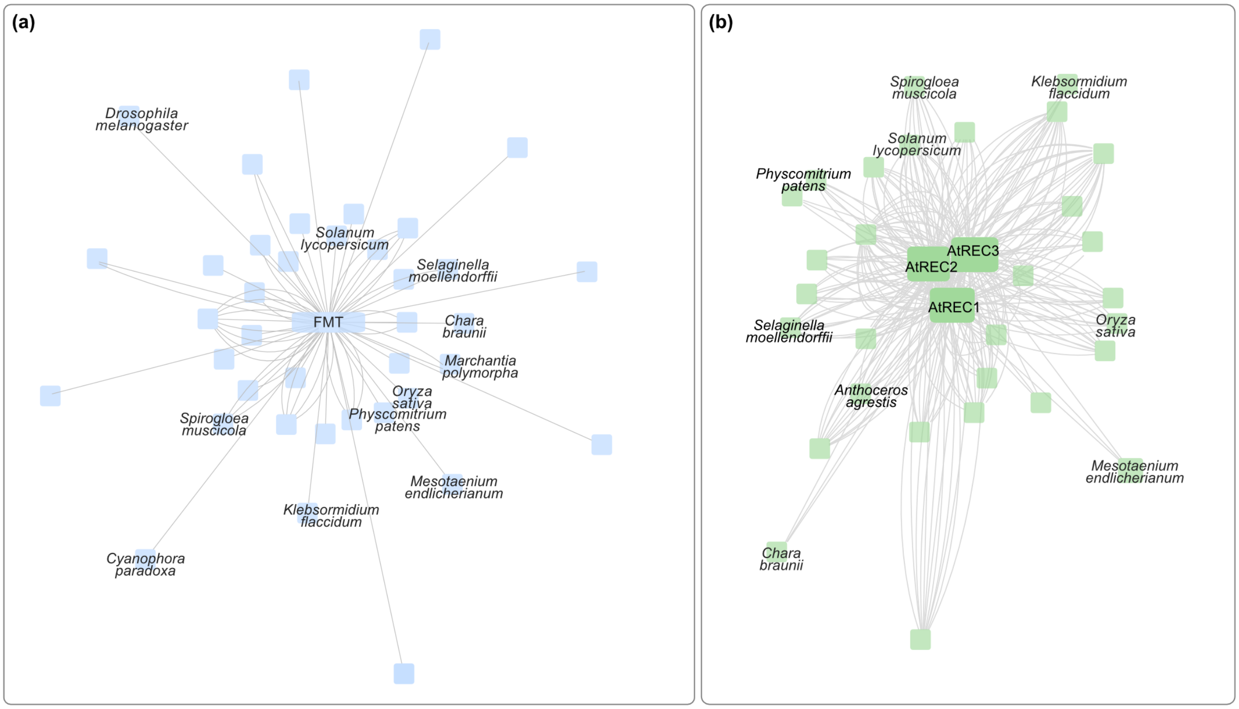


**Fig. S3: Mitochondrial and plastidial CLU family proteins form distinct clusters. (a)** Clusters of mitochondrial CLU family orthologues, resulting from whole genome clustering of 136 eukaryotes and annotated based on presence of Arabidopsis FRIENDLY (FMT, AT3G52140) and **(b)** of plastidial CLU family proteins, annotated based on the presence of the three Arabidopsis REC1, REC2 and REC3 proteins (AT1G01320, AT4G28080, AT1G15290). Each box represents a homologue; a subset of species (representing the evolutionary diversity of plants and animals) are labelled. Edges represent sequence similarities.

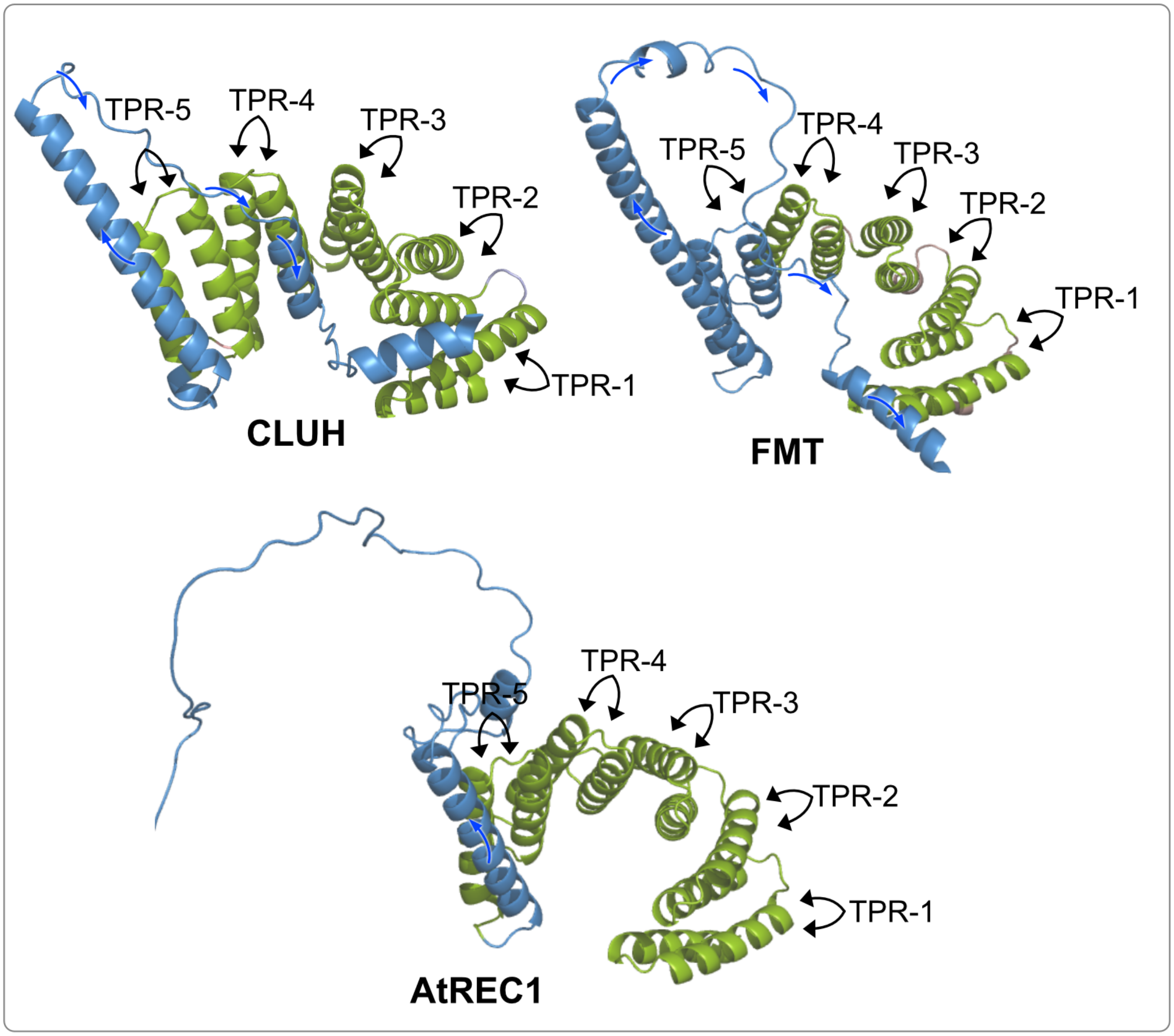


**Fig. S4: Structural differences between TPR regions of mitochondrial and plastidial CLU family proteins.** Structural predictions of the TPR regions from human and Arabidopsis mitochondrial CLU proteins (CLUH and FMT protein) suggest that an alpha helix after the fifth TPR motif of mitochondrial CLU proteins (shown in light blue) returns back (shown by cyan arrows) to the TPR region. In contrast, the same region in the plastidial homologue (AtREC) turns towards the opposite direction (shown by black arrows). Each unit TPR motif consists of a helix-turn-helix.

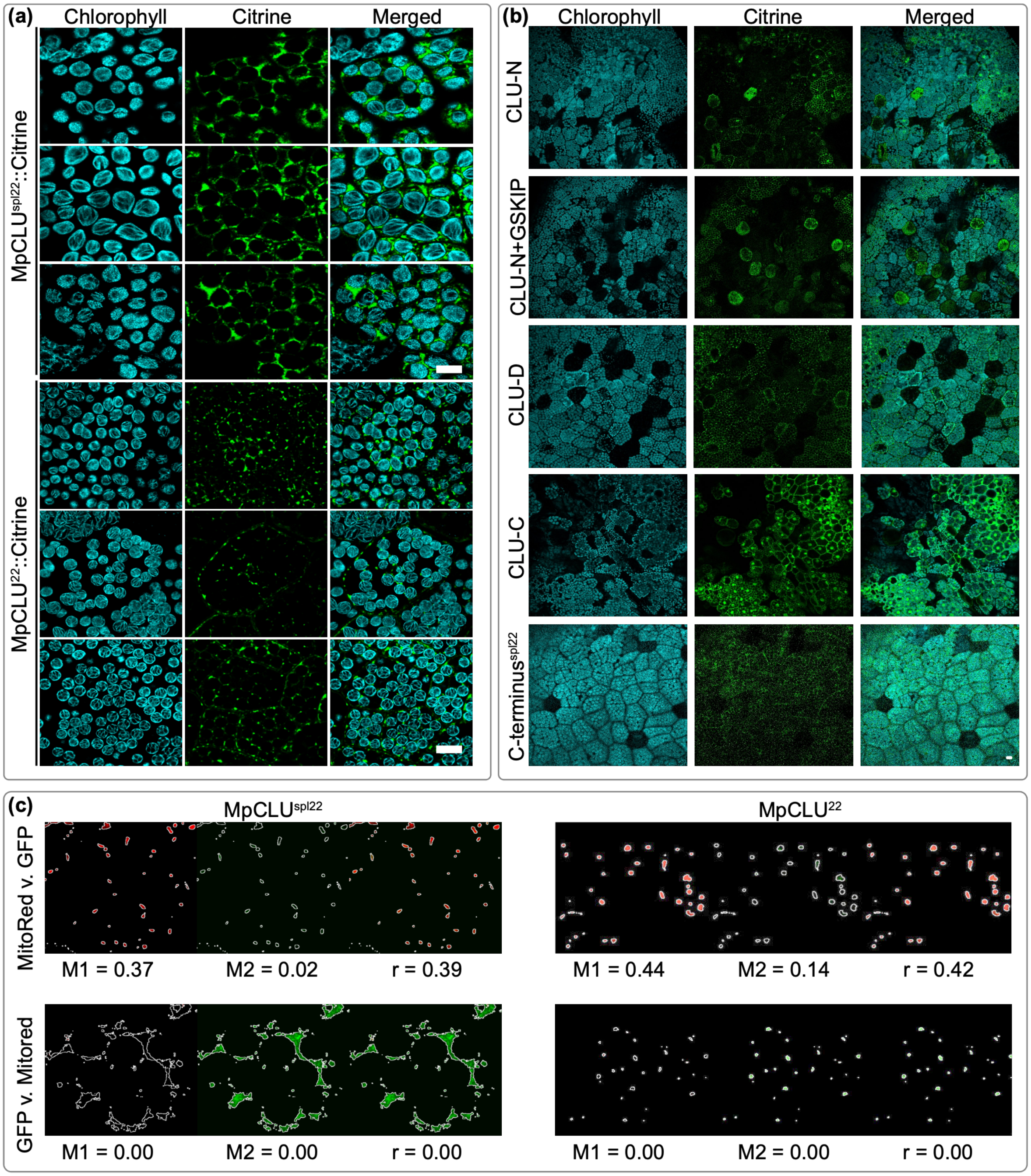


**Fig. S5: Additional subcellular localization of Mp*CLU*$^{spl22}$ and Mp*CLU*$^{22}$.** Confocal laser scanning images of **(a)** Mp*CLU*$^{22}$:Citrine and Mp*CLU*$^{spl22}$:Citrine, and **(b)** CLU-N, CLU-N + GSKIP, CLU-D, CLU-C, C-terminus$^{spl22}$ domains under 35S promoter in one-day-old thalli (chloroplasts are imaged through their autofluorescence). Scale bar: 10 µm. **(c)** Colocalization analyses for MpCLU$^{spl22}$ and MpCLU$^{22}$ constructs shows the GFP signal co-localises with the mitochondrial signal. Mander's coefficients M1 and M2, respectively, show a fraction of the reporter signal overlapping with mitochondria and vice versa (e. g. M1 = 0.37 and M2 = 0.02; i.e 37% of the mitochondrial region correlates with the GFP signal). R shows the correlation coefficient for intensities of the two channels.

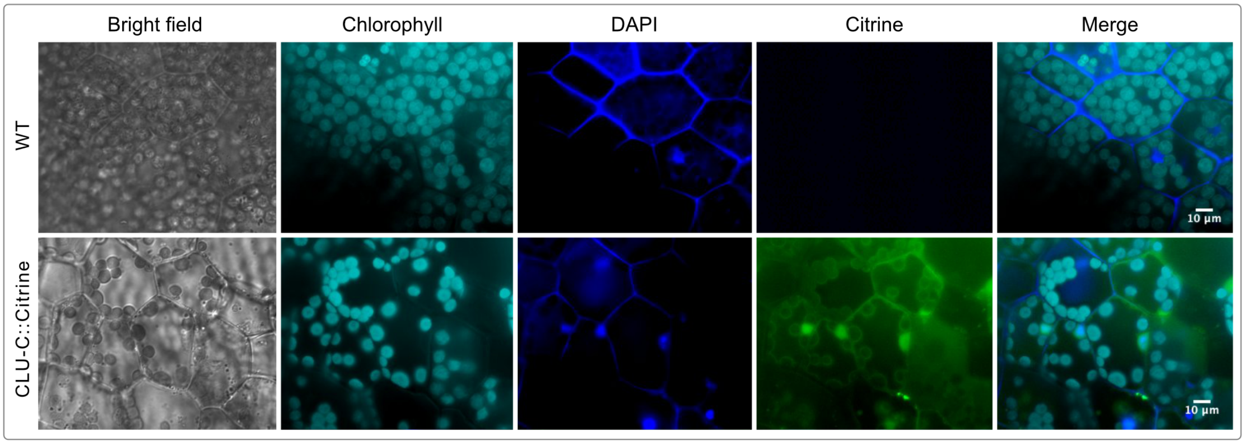


**Fig. S6: Nuclear localization of CLU-C.** Microscopy images of CLU-C:Citrine expression in one-day-old thalli stained with DAPI. Scale bars: 10 µm.

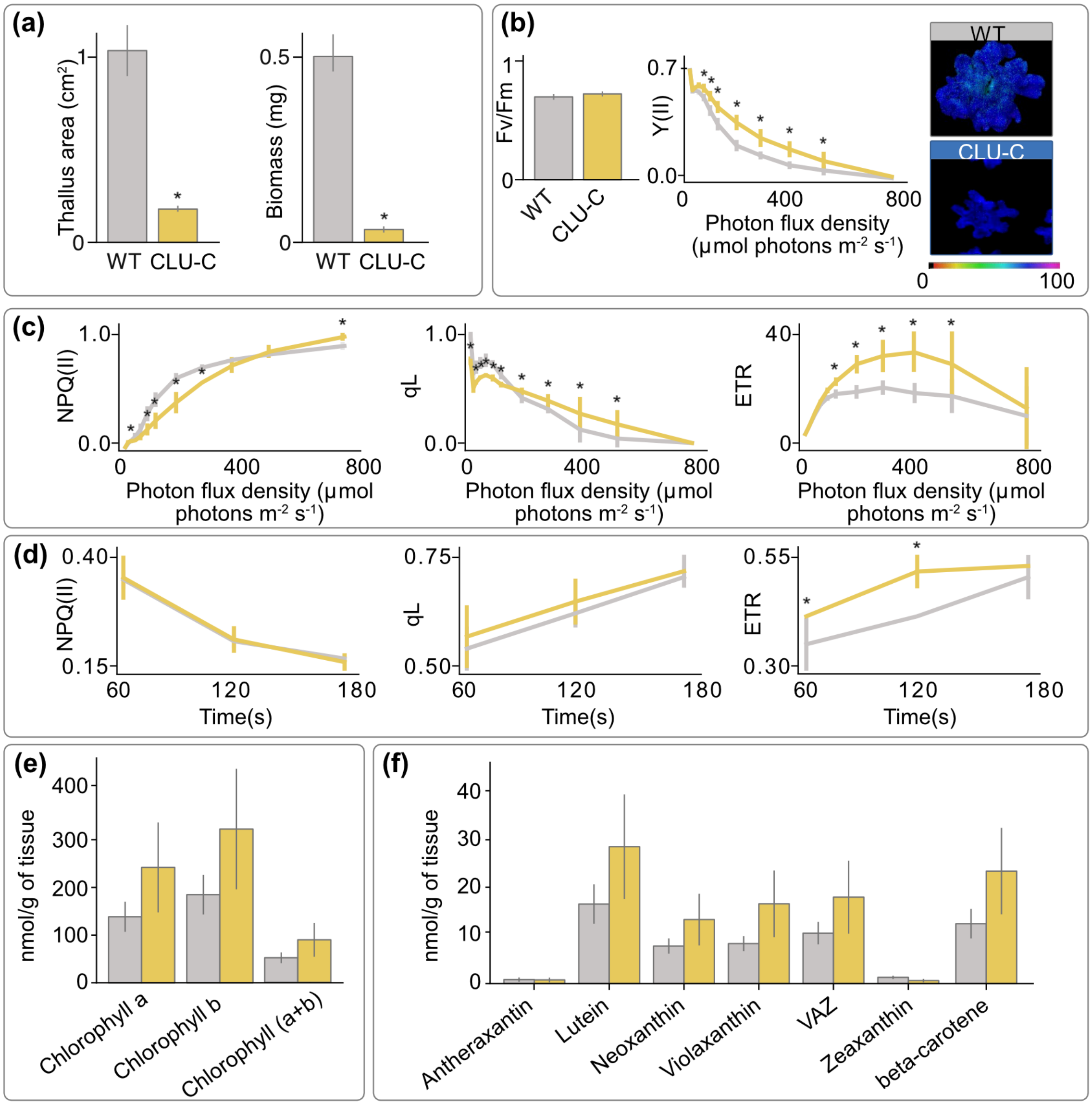


**Fig. S7: Effect of CLU-C:Citrine on photosynthesis activity and pigment accumulation. (a)**. Thallus area and biomass measurement of 14-day-old WT and CLU-C:Citrine plants (n=10; asterisks indicate a significant difference compared to the wild-type with p-value<0.05). **(b)** $F_v/F_m$ and Y(II) quantification of 14-day-old WT and CLU-C:Citrine plants (n=5) via an Imaging-PAM. **(c)** Quantification of the chlorophyll fluorescence parameters of WT (grey) and CLU-C:Citrine (yellow) plants with increasing photon flux density. **(d)** Quantification of the chlorophyll fluorescence parameters of WT and CLU-C:Citrine plants after PSII saturation with a constant photon flux density of 2 µmol photons $m^{-2}$ $s^{-1}$. **(e)** Chlorophyll and **(f)** carotenoids content for WT and CLU-C:Citrine plants (n=5). Asterisks indicate significant differences compared to the wild-type (p-value<0.05).

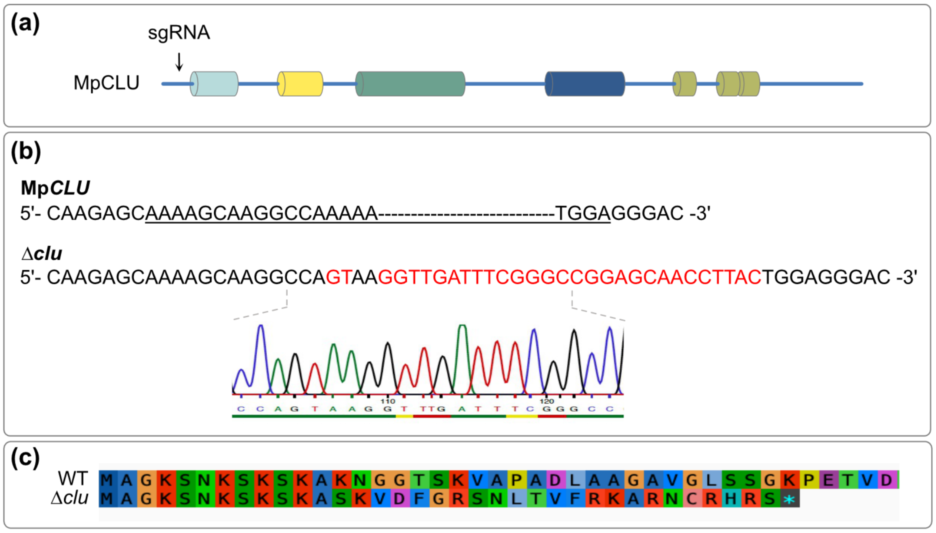


**Fig. S8: sgRNA design and sequencing results. (a)** Schematic of the MpCLU domain architecture and its conserved domains with the location of the guide RNA in the upstream region of the gene indicated. **(b)** Sequences of WT and Δ*clu*, with the mutations highlighted in red and the target site of the guide RNA underlined. Sequencing chromatogram of Δ*clu* below. **(c)** Amino acid sequence alignment of the WT and Δ*clu* sequence with the introduced stop codon at position 36.

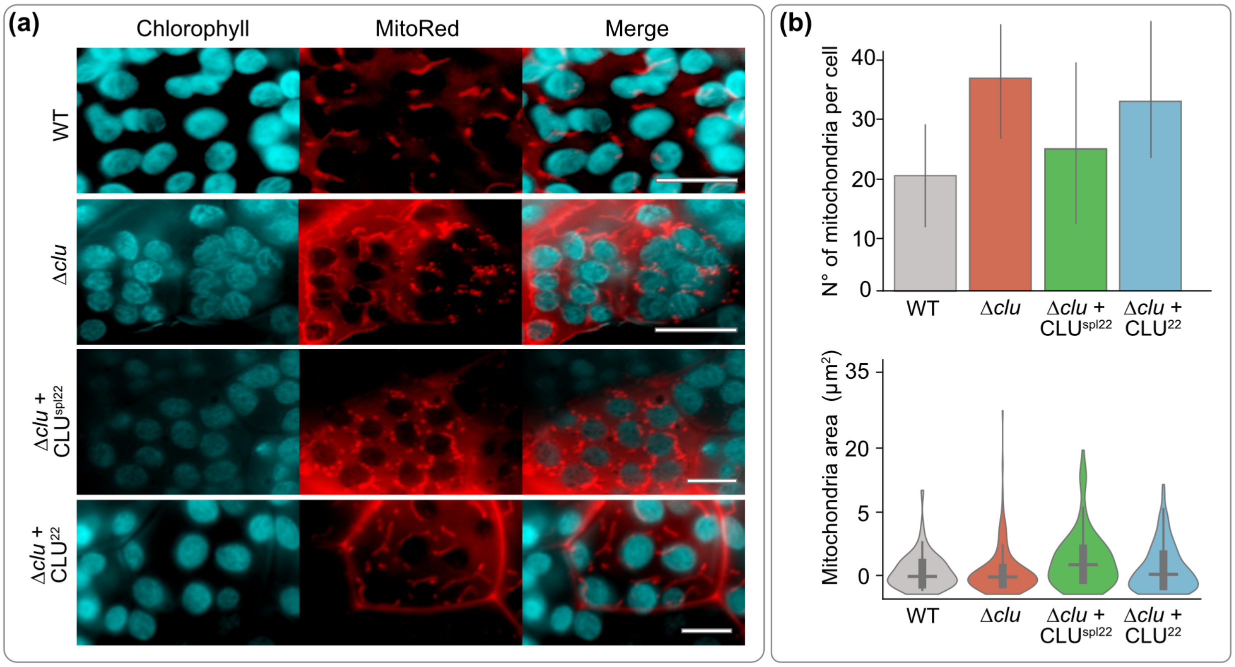


**Fig. S9: Loss-of-function of Mp*CLU* alters mitochondrial dynamics. (a)** Microscopy images of WT, Δ*clu*, Δ*clu* + MpCLU$^{spl22}$ and Δ*clu* + MpCLU$^{22}$ gemmae showing mitochondrial distribution. Mitochondria were stained with MitoRed, chloroplasts are shown through their chlorophyll autofluorescence. Scale bar: 10 µm. **(b)** Number of mitochondria per cell and total mitochondrial area in the cells of WT, Δ*clu*, Δ*clu* + MpCLU$^{spl22}$ and Δ*clu* + MpCLU$^{22}$. Asterisks indicate significant differences compared to the wild-type (p-value<0.05).

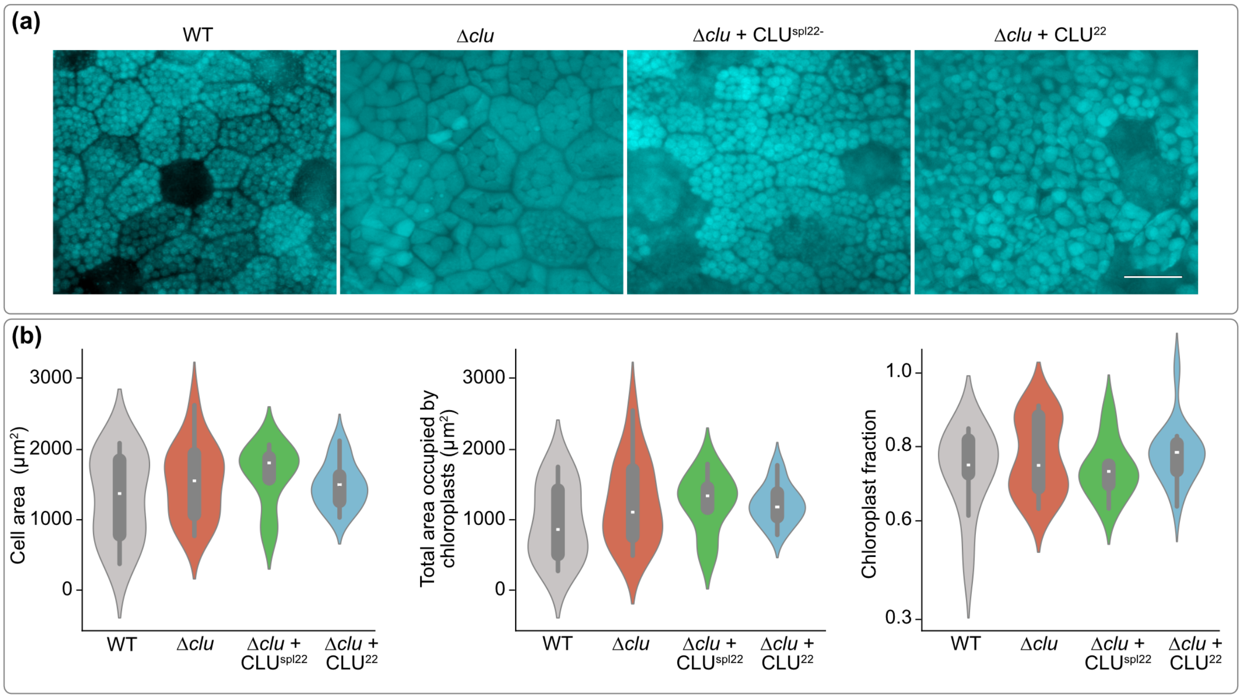


**Fig. S10: Loss of function of Mp*CLU* alters chloroplast morphology. (a)** Microscopy images of WT, Δ*clu*, Δ*clu* + MpCLU$^{spl22}$ and Δ*clu* + MpCLU$^{22}$ gemmae showing the chloroplast autofluorescence. **(b)** Cell area, total area occupied by chloroplast per cell, fraction of area occupied by chloroplast (total area occupied by chloroplast divided by area per cell) of WT, Δ*clu*, Δ*clu* + MpCLU$^{spl22}$ and Δ*clu* + MpCLU$^{22}$, n=10 for cell area; n=250 for chloroplast area. Asterisks indicate significant differences compared to the wild-type (p-value<0.05).

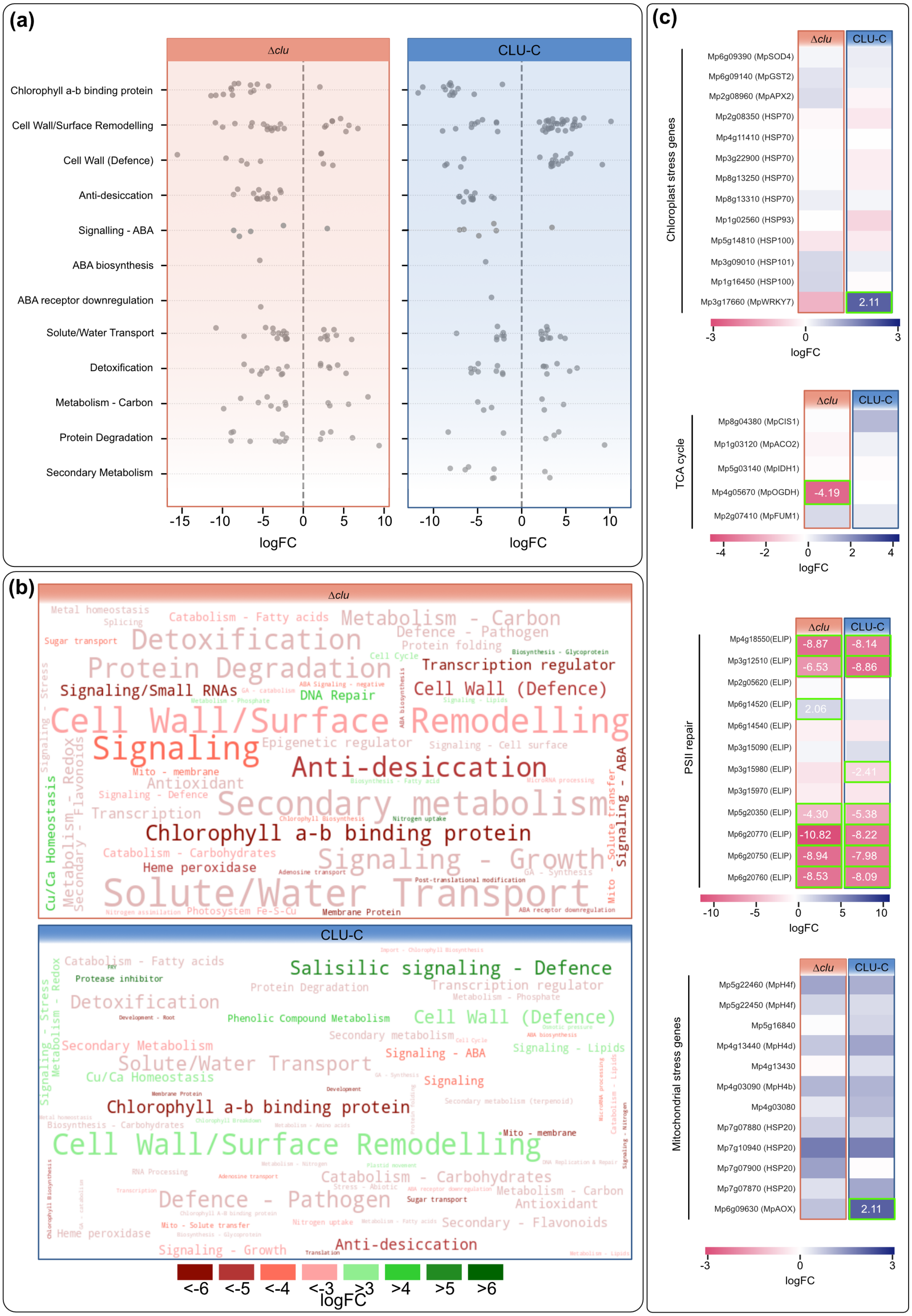

(a)
Δclu
CLU-C
Chlorophyll a-b binding protein
Cell Wall/Surface Remodelling
Cell Wall (Defence)
Anti-desiccation
Signalling - ABA
ABA biosynthesis
ABA receptor downregulation
Solute/Water Transport
Detoxification
Metabolism - Carbon
Protein Degradation
Secondary Metabolism
-15 -10 -5 0 5 10
-10 -5 0 5 10
logFC
logFC
(b)
Δclu
Metal homeostasis
Catabolism - Fatty acids
Metabolism - Carbon
Sugar transport
Detoxification
Defence - Pathogen
Protein folding
Protein Degradation
Transcription regulator
Signaling/Small RNAs
DNA Repair
Cell Wall (Defence)
Cell Wall/Surface Remodelling
Signaling
Epigenetic regulator
Signaling - Cell surface
Anti-desiccation
Mito - membrane
Antioxidant
Signaling - Defence
Secondary metabolism
Transcription
Chlorophyll a-b binding protein
Catabolism - Carbohydrates
Signaling - Growth
Heme peroxidase
Solute/Water Transport
Cu/Ca Homeostasis
Signaling - ABA
Photosystem Fe-S-Cu
Membrane Protein
CLU-C
Catabolism - Fatty acids
Salisilic signaling - Defence
Protease inhibitor
Protein Degradation
Transcription regulator
Detoxification
Phenolic Compound Metabolism
Cell Wall (Defence)
Secondary metabolism
Secondary Metabolism
Signaling - ABA
Signaling - Lipids
Solute/Water Transport
Signaling
Cu/Ca Homeostasis
Chlorophyll a-b binding protein
Mito - membrane
Cell Wall/Surface Remodelling
Catabolism - Carbohydrates
Defence - Pathogen
Metabolism - Carbon
Antioxidant
Sugar transport
Heme peroxidase
Secondary - Flavonoids
Signaling - Growth
Anti-desiccation
<-6 <-5 <-4 <-3 >3 >4 >5 >6
logFC
(c)
Chloroplast stress genes
Δclu
CLU-C
Mp6g09390 (MpSOD4)
Mp6g09140 (MpGST2)
Mp2g08960 (MpAPX2)
Mp2g08350 (HSP70)
Mp4g11410 (HSP70)
Mp3g22900 (HSP70)
Mp8g13250 (HSP70)
Mp8g13310 (HSP70)
Mp1g02560 (HSP93)
Mp5g14810 (HSP100)
Mp3g09010 (HSP101)
Mp1g16450 (HSP100)
Mp3g17660 (MpWRKY7)
2.11
-3 0 3
logFC
TCA cycle
Δclu
CLU-C
Mp8g04380 (MpCIS1)
Mp1g03120 (MpACO2)
Mp5g03140 (MpIDH1)
Mp4g05670 (MpOGDH)
-4.19
Mp2g07410 (MpFUM1)
-4 -2 0 2 4
logFC
PSII repair
Δclu
CLU-C
Mp4g18550(ELIP) -8.87 -8.14
Mp3g12510 (ELIP) -6.53 -8.86
Mp2g05620 (ELIP)
Mp6g14520 (ELIP) 2.06
Mp6g14540 (ELIP)
Mp3g15090 (ELIP)
Mp3g15980 (ELIP) -2.41
Mp3g15970 (ELIP)
Mp5g20350 (ELIP) -4.30 -5.38
Mp6g20770 (ELIP) -10.82 -8.22
Mp6g20750 (ELIP) -8.94 -7.98
Mp6g20760 (ELIP) -8.53 -8.09
-10 -5 0 5 10
logFC
Mitochondrial stress genes
Δclu
CLU-C
Mp5g22460 (MpH4f)
Mp5g22450 (MpH4f)
Mp5g16840
Mp4g13440 (MpH4d)
Mp4g13430
Mp4g03090 (MpH4b)
Mp4g03080
Mp7g07880 (HSP20)
Mp7g10940 (HSP20)
Mp7g07900 (HSP20)
Mp7g07870 (HSP20)
Mp6g09630 (MpAOX)
2.11
-3 0 3
logFC

**Fig. S11: Functional categories of differentially expressed genes (DEGs) in Δ*clu* and CLU-C::Citrine lines. (a)** Each dot represents a gene differentially expressed (logFC>2) in Δ*clu* and CLU-C lines, across prominent functional categories shown on the left. **(b)** Word clouds of the functional categories of DEGs in Δ*clu* (up) and CLU-C::Citrine lines (down), where the size of the words indicate the total number of DEGs under that category (i.e. the bigger the word, the greater the number of DEGs in that category). The color of the categories indicates median logFC across all genes in that category as per the key on the bottom right (e.g. categories labelled in the darkest shade of green and logFC=6 suggest that a typical gene under that category was heavily up regulated). **(c)** Expression profiles of individual genes from functional categories of interest plotted as a heatmap; statistically significant differences with logFC>2 are marked by green borders and with the expression value provided.

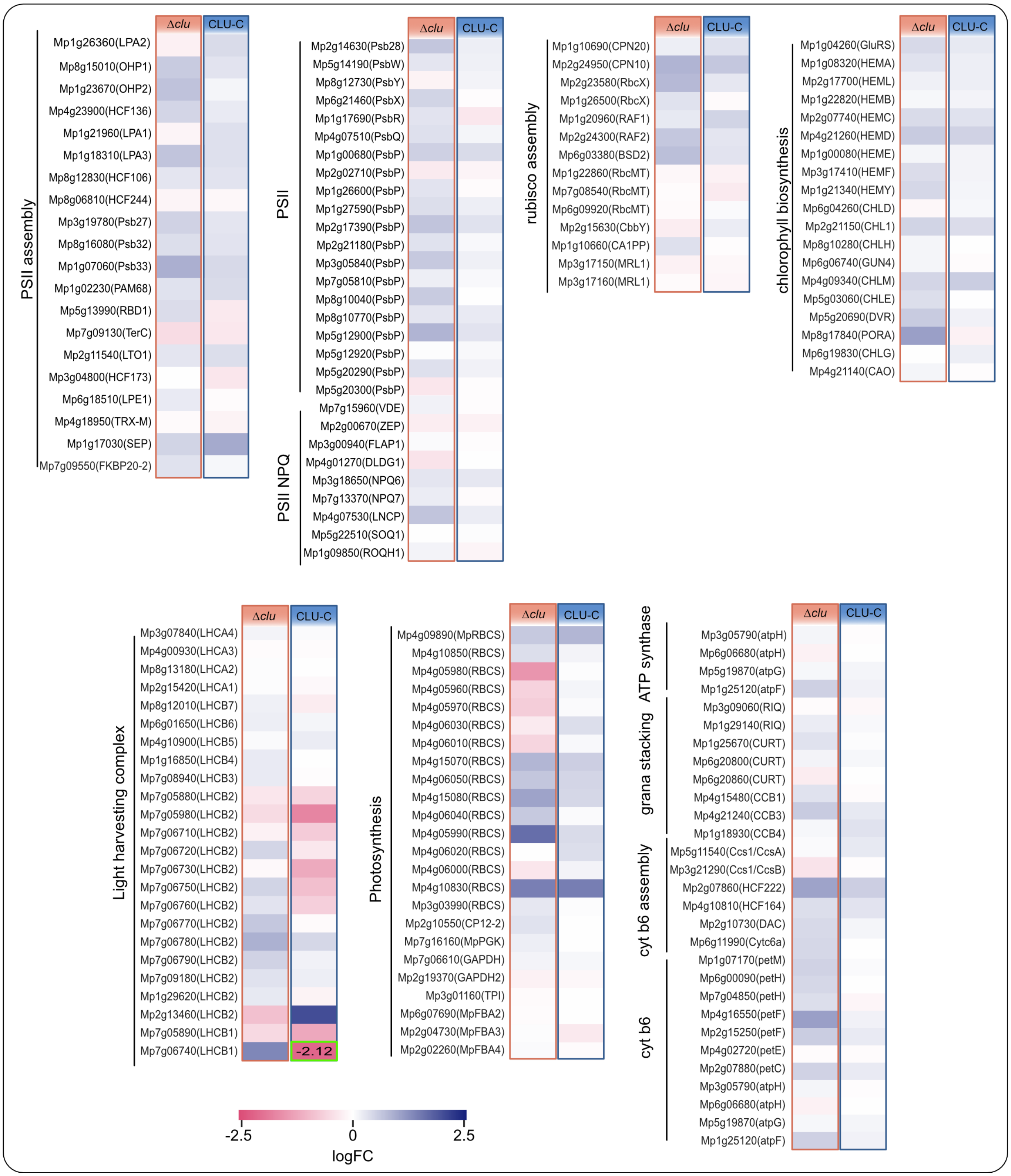


**Fig. S12: Global gene expression changes in Δ*clu* and CLU-C:Citrine plants.** Expression profiles of individual genes from functional categories of interest plotted as a heatmap; statistically significant differences with logFC>2 are marked by green borders and a provided expression value.

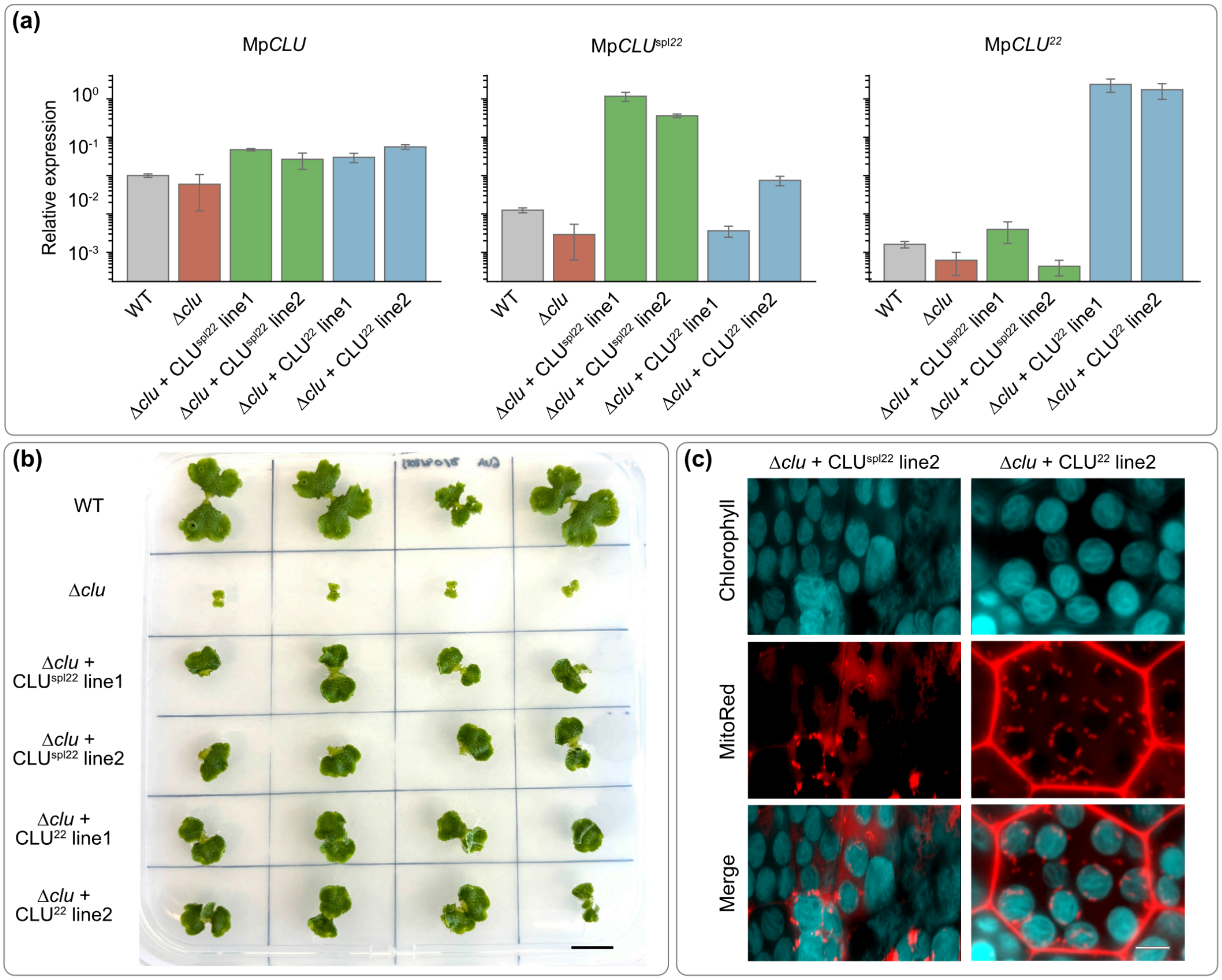


**Fig. S13:** Additional Δ*clu* + MpCLU$^{spl22}$ and Δ*clu* + MpCLU$^{22}$ lines. **(a)** Expression levels of CLU splicing variants, relative to that of actin, measured by RT-qPCR. **(b)** Representative images of WT, Δ*clu*, Δ*clu* + CLU$^{spl22}$ line 1 and Δ*clu* + CLU$^{22}$ line 2 plants grown from gemmae till day 14 (scale bar: 1 cm). **(c)** One-day-old thalli pictures of WT, Δ*clu* and Δ*clu* rescued with either CLU$^{spl22}$ or CLU$^{22}$. Plastids imaged through their autofluorescence (a-b) and mitochondria through MitoTracker Red. Scale bar: 10 µm.

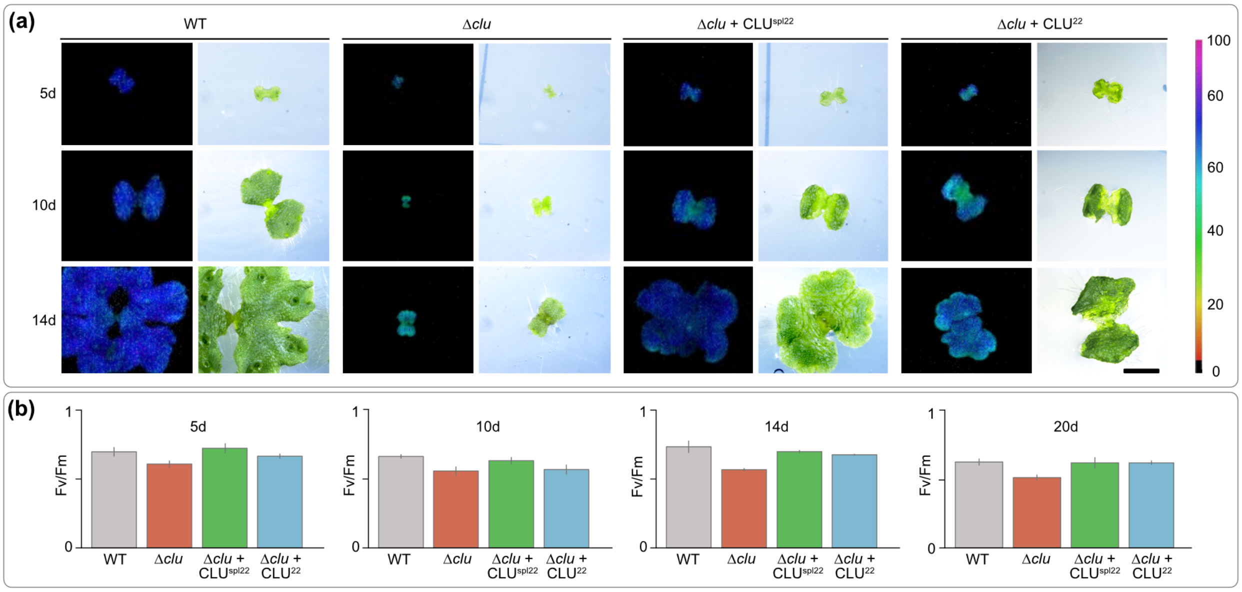


**Fig. S14: Effect on photosynthesis activity of Mp*CLU*. (a)** Representative images of thalli and the chlorophyll fluorescence parameter $F_v/F_m$ of WT, Δ*clu*, Δ*clu* + MpCLU$^{spl22}$ and Δ*clu* + MpCLU$^{22}$. Scale bar: 0,2 cm. Data is represented as means +/- SD; n = 5. **(b)** Quantification of the chlorophyll fluorescence parameter $F_v/F_m$ of WT, Δ*clu*, Δ*clu* + MpCLU$^{spl22}$ and Δ*clu* + MpCLU$^{22}$ (n=5).

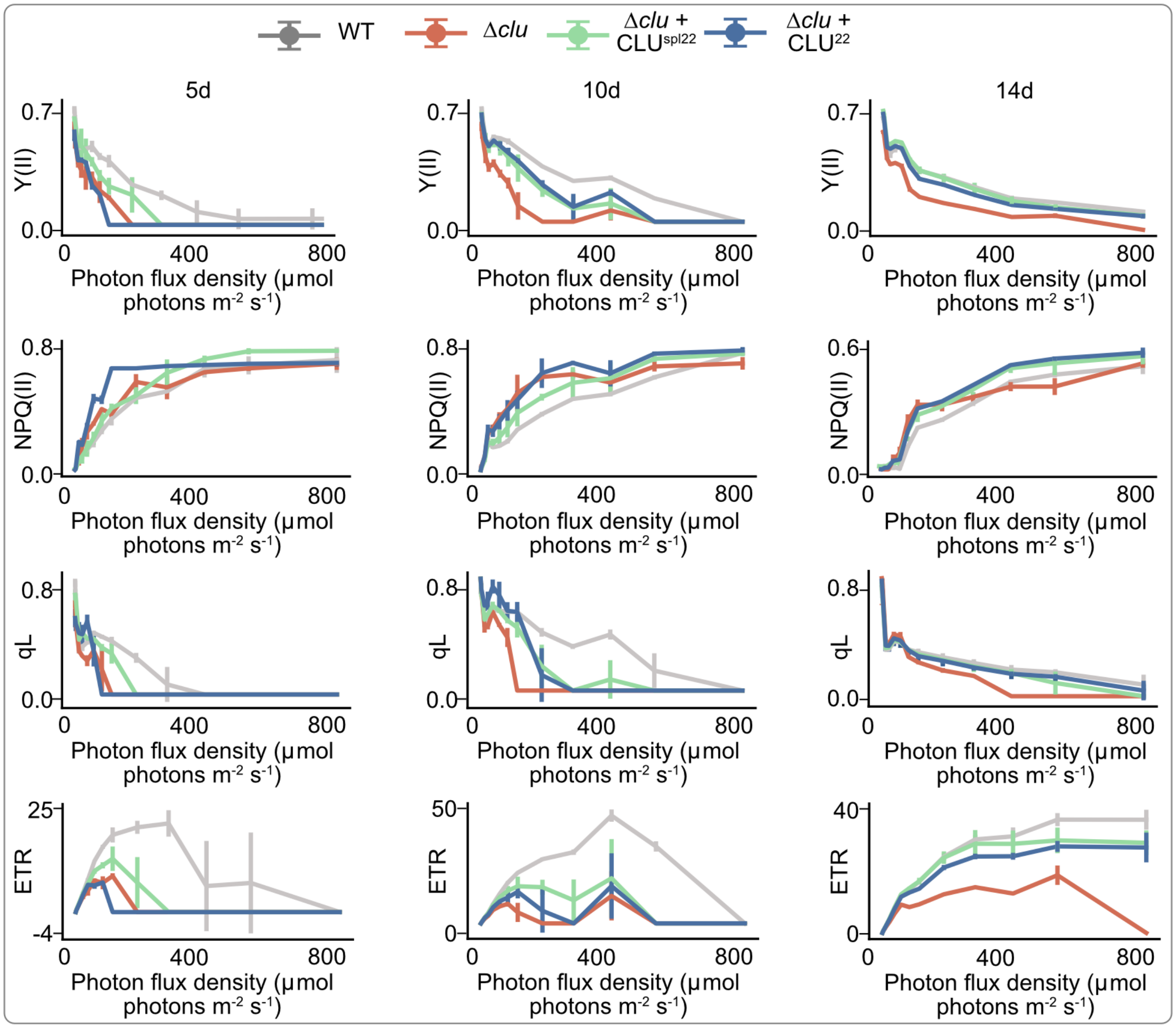


**Fig. S15: Effect of Mp*CLU* on photosynthesis activity.** Quantification of the chlorophyll fluorescence parameters of WT, Δ*clu*, Δ*clu* + MpCLU$^{spl22}$ and Δ*clu* + MpCLU$^{22}$ with increasing photon flux density, on 5-, 10-, and 14-day-old plants (n=5).

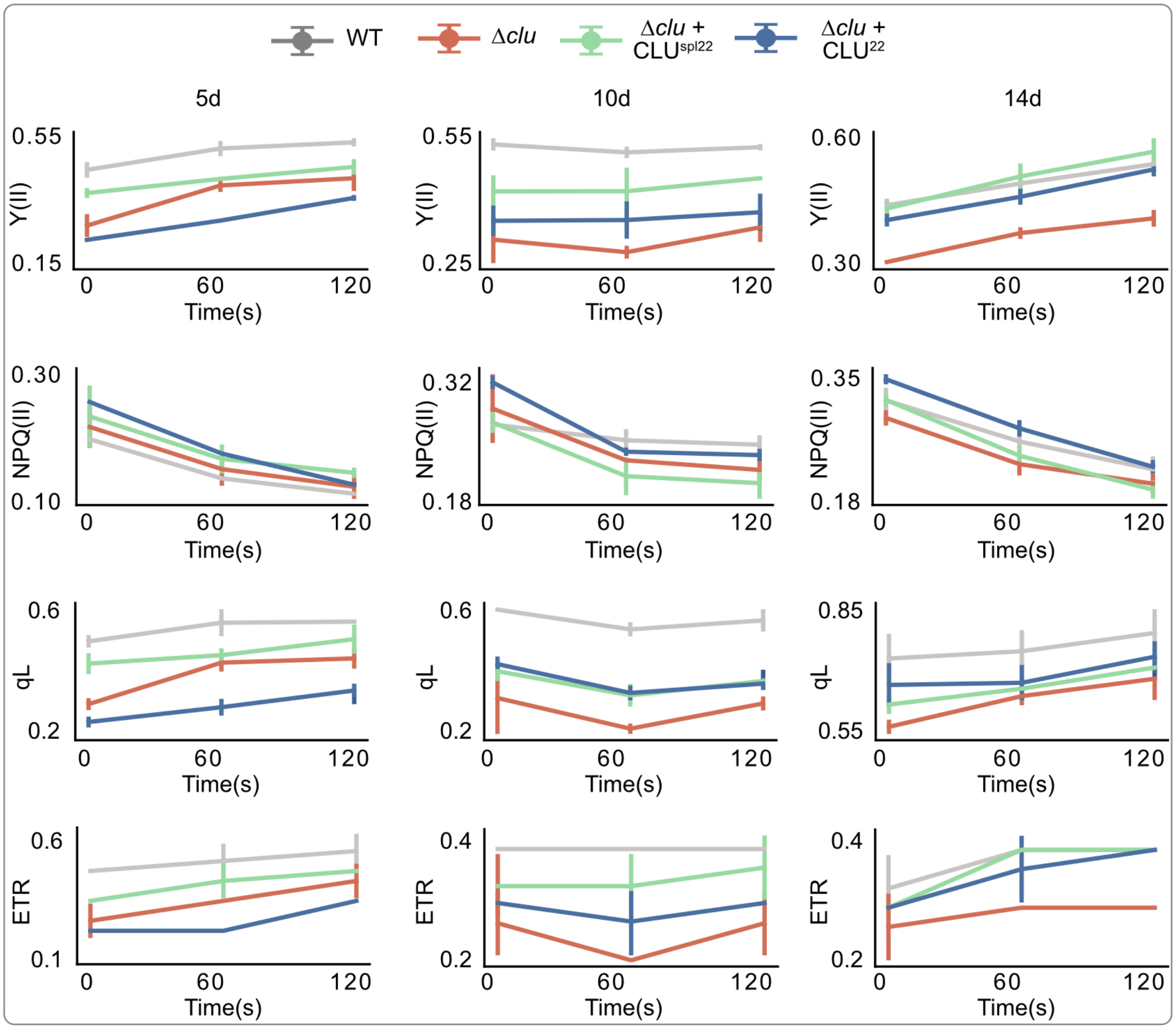


**Fig. S16: Effect of Mp*CLU* on PSII recovery after saturation.** Quantification of the chlorophyll fluorescence parameters of WT, Δ*clu*, Δ*clu* + $MpCLU^{spl22}$ and Δ*clu* + $MpCLU^{22}$ with constant photon flux density of 2 µmol photons $m^{-2}$ $s^{-1}$ after PSII was saturated on 5-, 10-, and 14 days old plants (n=5). Asterisks indicate significant differences compared to the wild-type (p-value<0.05).
